# Carbohydrate binding domains facilitate efficient oligosaccharides synthesis by enhancing mutant catalytic domain transglycosylation activity

**DOI:** 10.1101/2020.05.06.081356

**Authors:** Chandra Kanth Bandi, Antonio Goncalves, Sai Venkatesh Pingali, Shishir P. S. Chundawat

**Author notes:** **Corresponding Author Name:** Shishir P. S. Chundawat (ORCID: 0000-0003-3677-6735) **Corresponding Author Email:**.

## Abstract

Chemoenzymatic approaches using carbohydrate-active enzymes (CAZymes) offer a promising avenue for synthesis of glycans like oligosaccharides. Here, we report a novel chemoenzymatic route for cellodextrins synthesis employed by chimeric CAZymes, akin to native glycosyltransferases, involving the unprecedented participation of a ‘non-catalytic’ lectin-like or carbohydrate-binding domains (CBMs) in the catalytic step for glycosidic bond synthesis using β-cellobiosyl donor sugars as activated substrates. CBMs are often thought to play a passive substrate targeting role in enzymatic glycosylation reactions mostly via overcoming substrate diffusion limitations for tethered catalytic domains (CDs) but are not known to participate directly in any nucleophilic substitution mechanisms that impact the actual glycosyl transfer step. Our study provides evidence for the direct participation of CBMs in the catalytic reaction step for β-glucan glycosidic bonds synthesis enhancing activity for CBM-based CAZyme chimeras by >140-fold over CDs alone. Dynamic intra-domain interactions that facilitate this poorly understood reaction mechanism were further revealed by small-angle X-ray scattering structural analysis along with detailed mutagenesis studies to shed light on our current limited understanding of similar transglycosylation-type reaction mechanisms. In summary, our study provides a novel strategy for engineering similar CBM-based CAZyme chimeras for synthesis of bespoke oligosaccharides using simple activated sugar monomers.

## Introduction

Glycans are complex biomacromolecules (e.g., cellulose and N-linked glycans) composed of monosaccharides linked via glycosidic linkages to either other carbohydrates, lipids, or proteins to form a combinatorially large number of biomolecular structures that play still poorly understood structural and functional roles in biological systems (Moon et al., 2011; Varki and Lowe, 2009). There has been a tremendous effort over the last decade to advance the broad field of glycosciences via development of novel methods for synthesis or modification of bespoke glycans (National Research Council, 2012). Hybrid chemoenzymatic approaches using engineered glycosidases or glycosyl hydrolases (GHs) offer a promising avenue for synthesis of glycans like oligosaccharides (Wang et al., 2013b). GHs are more abundantly available in genomic databases, are often well characterized, express readily using *E. coli*, and have promiscuous substrate specificity that make them more attractive biocatalysts compared to traditional catalytic synthesis methods (Kiessling and Splain, 2010; Pardo-Vargas et al., 2018; Plante et al., 2001; Yu et al., 2019) or natural glycan synthesizing enzymes such as glycosyl transferases (GTs) (Boltje et al., 2009; Lairson et al., 2008; Wang and Davis, 2013; Wang and Lomino, 2012). GHs are grouped into various families based on amino-acid sequence similarity (Henrissat and Davies, 1997), currently numbering at 166+ families as curated on the Carbohydrate-Active enZyme (CAZyme) database (Cantarel et al., 2009; Lombard et al., 2014), and still growing with newly sequenced genomes and metagenomes becoming readily available. CAZymes like GHs or GTs are further classified either as retaining or inverting type, depending on whether the stereochemistry at the anomeric carbon for the product (versus the reactant) is either retained or inverted (Davies and Henrissat, 1995), respectively. As first proposed by Koshland for glycosidases (Koshland, 1953), retaining enzymes require a classical two-step double displacement hydrolysis mechanism (i.e., S_N_2) with a glycosyl-enzyme intermediate (GEI) formed via a covalent bond between the cleaved substrate and the protein in the alternate orientation (e.g., GEI for retaining enzyme will have an α-bond, as opposed to the β-orientation of the reactant and product). While most native glycosidases catalyze the hydrolysis of glycosidic linkages, many GHs can also produce oligosaccharides following a competing transglycosylation mechanism if a suitably localized glycosyl acceptor group is present instead of a water molecule within the enzyme active site (Bissaro et al., 2015). Retaining GHs that show significant transglycosylation reactivity are also thought to utilize a similar S_N_2 type general mechanism, except that rather than nucleophilic attack by water, the attack is instigated by a hydroxyl oxygen on an acceptor sugar placed adjacent to the GEI complex (Figure 1).

**Figure 1.**
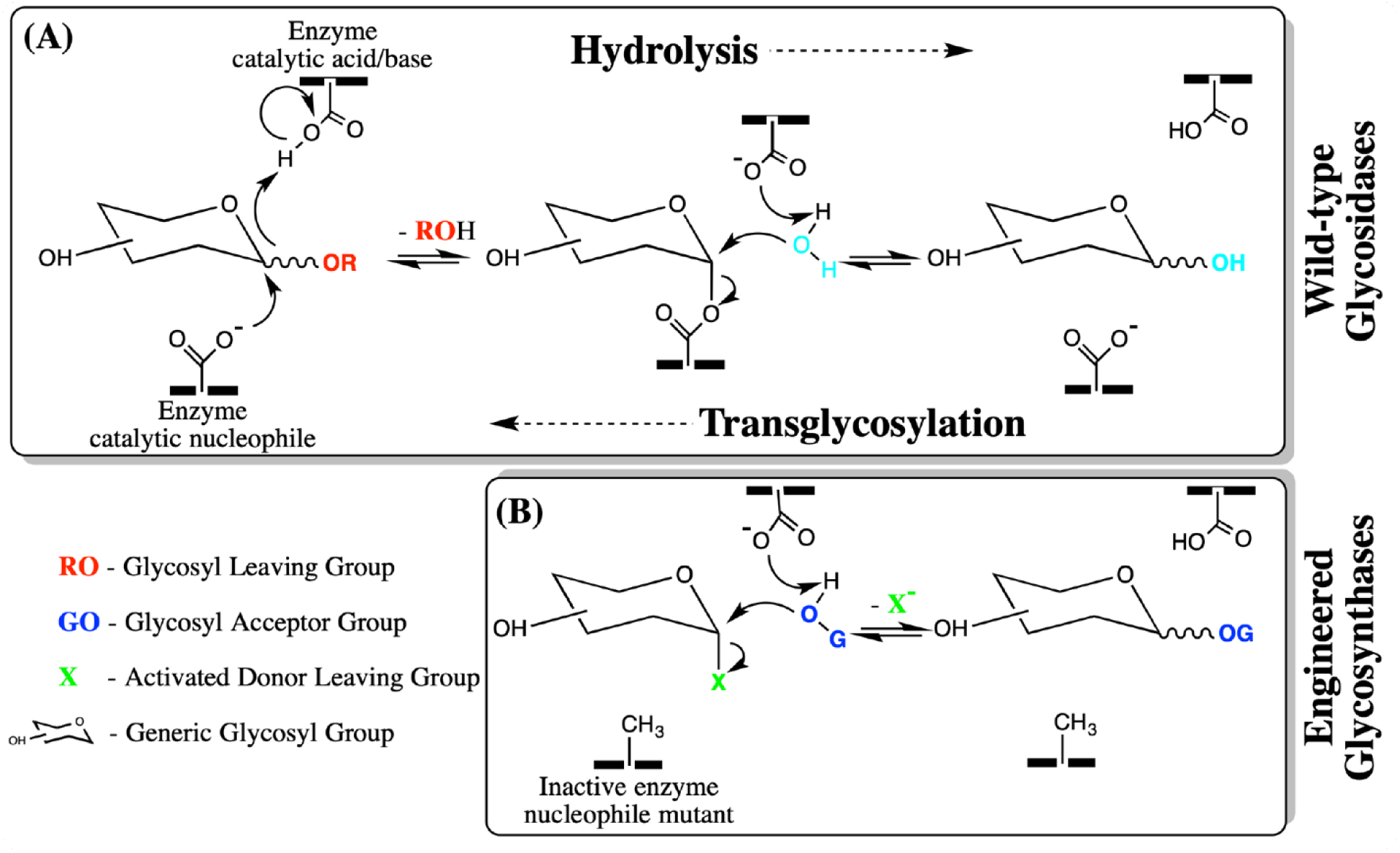
Overview of generic S_N_2 based two-step mechanism employed by retaining glycosyl hydrolases and corresponding nucleophile mutants (e.g., glycosynthases) to facilitate both hydrolysis and transglycosylation type reactions. Here, (A) depicts the glycosylation step initiated by the enzyme catalytic nucleophile residue that results in the formation of a Glucosyl-Enzyme Intermediate (GEI). In the presence of water acting as a nucleophile, hydrolysis (and subsequent deglycosylation of the GEI) takes place for most native wild-type glycosidases. In the presence of a suitable glycosyl acceptor group, transglycosylation can take place as well for some native or engineered glycosidases. (B) Mutations at the nucleophile site to develop engineered transglycosidases mutants, like glycosynthases, can often facilitate efficient synthesis of glycans like oligosaccharides using simple activated donor sugars that mimic the GEI conformation.

Unfortunately, transglycosylation reactions for most native GHs often suffer from low product yields since the transglycosylation product is also the native substrate for GHs (Figure 1A). Mutant glycosidases, like glycosynthases (GSs), offer a simple solution to address this issue to synthesize glycans (Cobucci-Ponzano et al., 2011; Dalia Shallom et al., 2004; David L. Zechel et al., 2003; Mayes et al., 2016; Payne et al., 2015). The wild-type GH nucleophilic residue such as aspartate or glutamate is often mutated to one that can no longer accept a proton (e.g., alanine, glycine or serine) (Cobucci-Ponzano and Moracci, 2012). These mutant transglycosidases or GSs perform glycan synthesis in the presence of activated donor sugars whose structures mimic the GEI. Like the native GEI, these donor sugars have the opposite stereochemistry at the anomeric center from the native reactant, and the anomeric carbon is bonded to a good electrophilic leaving group (Cobucci-Ponzano and Moracci, 2012). Nucleophilic-site mutations introduced into retaining GHs to create more efficient transglycosidases like GSs were thought originally to not impact the established S_N_2 reaction mechanism (Figure 1B). However, a recent study has challenged this paradigm showing that a nucleophile mutant of a GH family 1 β-glycosidase follows an S_N_i-like mechanism to synthesize β-glycosides in the presence of activated β-donors like p-nitrophenyl (pNP) based glycosides (Iglesias-Fernández et al., 2017). Currently there are no other reports in the literature of β-retaining enzymes that follow a front-facing S_N_i-like mechanism. It is also unclear if other CAZyme domains found often associated with GHs would impact such a front-facing reaction mechanism in engineered β-retaining GH enzymes. This would be analogous to the role of lectin-like domains identified in the case of multidomain GTs that perform the glycosyl transfer step with nucleotide donor sugars using a classical S_N_i type mechanism (Lira-Navarrete et al., 2015).

GHs are often found naturally associated with non-catalytic auxiliary domains, like CBMs, that specifically recognize and bind to carbohydrates (Boraston et al., 2004; Van Tilbeurgh et al., 1986). CBMs are thought to increase the catalytic efficiency of CAZymes such as GHs, GTs, and carbohydrate esterases (CEs) by mostly overcoming substrate diffusion limitations. While removal of CBMs from full-length GHs has been shown to cause significant decrease in activity towards mostly larger polysaccharides (Burstein et al., 2009; Gal et al., 1997; Gaudin et al., 2000), CBMs are thought to not impact the tethered catalytic domain activity towards soluble substrates like model pNP-glycosides or shorter oligosaccharides (Tomme et al., 1988). CBMs particularly modulate the hydrolytic activity of GHs by disrupting or modifying the polysaccharide structure and increasing the effective substrate concentration near the catalytic domain (Boraston et al., 2004; Din et al., 1991). Although many studies have highlighted the influence of CBMs on the hydrolytic activity of GHs, the effect of CBMs on transglycosylation activity of GHs or GSs is still not well understood from a mechanistic point-of-view. For example, plant cell wall expansins and microbial GH family 45 enzymes are structurally related CBM-containing CAZyme families that show significant plant cell wall growth/extension and transglycosylation/hydrolysis activity, respectively (Wang et al., 2013a; Yennawar et al., 2006). Surprisingly, plant expansins lack a traditional catalytic base in the active site thought to be necessary for catalytic activity and operate via molecular mechanisms likely involving CBMs that are still not understood (Kerff et al., 2008). Such issues have broadly hampered research in the area of engineering better transgenic plants and consolidated bioprocess (CBP) cellulolytic microbes which ultimately has direct impact on plant biomass harvest yields and cellulosic biofuel production costs, respectively.

Here, we report the influence of CBMs on the transglycosylation activity of a β-retaining chimeric transglycosidase enzyme design for the synthesis of β-1,4-glucan oligosaccharides (i.e., cellodextrins) that surprisingly followed a poorly understood synthase mechanism even after mutation of the nucleophilic active site instead of the classical S_N_2-type mechanism seen for the native enzyme. The native GH scaffold chosen for this study belonged to a native β-retaining family 5 cellulase, called CelE from a well-known CBP microbe *Clostridium thermocellum*, with characteristic catalytic nucleophile and acid/base residues. Unsurprisingly, the nucleophile site mutation (to Alanine) of CelE’s CD domain alone abrogated enzyme activity on soluble substrates like pNP-β-D-cellobiose in line with the expected role of a true nucleophile residue on the catalytic turnover of a β-retaining enzyme following an S_N_2 type mechanism. However, fusion of these catalytically-inactive CelE nucleophile mutants to a *C. thermocellum* CBM3a domain associated with its native linker converted the inactive CD into an active transglycosidases that produced cellodextrin based glycosylation products. These studies also revealed, for the first-time, the unexpected involvement of CBMs and optimum linker domain in the actual catalytic reaction step of chimeric CAZymes. Through detailed biochemical characterization of several engineered enzyme constructs, along with complementary structural analyses, we also provide preliminary supporting evidence for previously unknown competing reaction mechanisms (e.g., S_N_i vs. S_N_2 mechanism) utilized by such CAZymes.

## Materials and Methods

See Supporting Information appendix (Supplementary Text) for all materials and methods relevant to this study.

## Results

### CBM3a recovers CelE nucleophile mutants transglycosylation and hydrolytic activity

CelE from *Clostridium* (*Ruminiclostridium) thermocellum* CBP microbe belongs to the glycosyl hydrolase family 5 (GH5) and is known to follow a retaining mechanism during hydrolysis of soluble cello-oligosaccharides and insoluble amorphous cellulose based polysaccharides to mostly cellobiose (Aspeborg et al., 2012; Gaudin et al., 2000; Walker et al., 2015). We were initially interested to test the transglycosidase activity of CelE catalytic domain alone for synthesis of β-glucans and therefore decided to introduce suitable mutations to reduce any competing hydrolysis activity. Retaining enzymes, like CelE from GH5 with a common (β/α)_8_ TIM barrel fold, have been successfully engineered into GSs in the past by mutating the native catalytic nucleophile into a small polar amino acid as first shown by Withers and co-workers (Mackenzie et al., 1998). In order to engineer CelE into a glycosynthase, the catalytic nucleophile site (E316), identified from the UniProt database (UniProt ID: P10477), was mutated to either an Alanine (E316A), Serine (E316S), or Glycine (E316G) residue. These mutant genes were next also fused to a *C. thermocellum* CBM3a domain linked by a 42aa flexible linker from the *C. thermocellum* scaffoldin protein CipA (Cthe_3077) on the C-terminus (Figure 2). The respective CelE and CelE-CBM3a gene constructs were cloned into an expression vector with N-terminal his-tags optimized for *E. coli* expression and the expressed enzyme purification was performed using standard metal affinity chromatography (Takasuka et al., 2014). The purified nucleophile mutant proteins were tested for recovered hydrolytic activity on amorphous cellulose (insoluble) and pNP-β-D-cellobiose (soluble) substrates in the presence of exogenously added chemical rescue agents such as azide and formate ions (**SI Figure S1**). The reactivation experiments revealed that hydrolytic activity for none of the nucleophilic mutants of CelE could be chemically rescued by either azide or formate. These results suggest that it would be likely not possible to form a true glycosynthase from CelE, based on previous reports that suggest a strong correlation between chemical rescue and glycosynthase activity (David L. Zechel et al., 2003). However, surprisingly, we observed that only the serine and glycine mutants of CelE-CBM3a nucleophile mutants, showed significant release of the pNP leaving group upon long periods of incubation with pNP-β-D-cellobiose (pNPC) as substrate (Figure 2A). An orcinol visualization stained thin-layer chromatographic (TLC) analysis of the reaction mixtures indicated the formation of cello-oligosaccharides with a degree of polymerization or DP≥3 (Figure 2B). The two dominant equilibrium reaction products were identified to be cellotriose (DP3) and cellotetraose (DP4) by comparing the TLC retention factors of the eluting compounds compared to their respective standards. The UV image of the TLC plate prior to orcinol staining also confirmed the presence of pNP containing higher molecular weight sugar polymers like pNP-cellotetraose (**SI Figure S2**). Interestingly, these oligomeric products synthesized by the E316G nucleophile mutants of CelE-CBM3a were subsequently found to be fully hydrolyzed upon the addition of wild type CelE enzyme (Figure 2C). This suggested that the glycosidic linkage formed is β(1,4) and the transglycosylation products formed can be recognized by the native CelE enzyme active site and fully hydrolyzed into cellobiose alone. Since, CelE belongs to very large family of multifunctional GH5 enzymes that predominantly catalyze hydrolysis of β(1,4) linked sugars, we expected the synthesized higher DP products to be β-retaining as well like pNPC substrates. Because of the multifunctional nature of these GH5 enzymes, we also tested activity of all enzyme constructs towards other glycosidic linkage based polysaccharides such as β(1,3) glucan and starch α(1,4). However, the native enzyme or its nucleophile mutants, either with or without the CBM3a domain, showed no activity towards these substrates (Figure 2D) implying that the higher DP products formed were not β(1,3) or α(1,4) linked gluco-oligosaccharides. These observations regarding the exact molecular weight and type of glycosidic linkage of the oligosaccharide products have been corroborated by liquid chromatography and mass spectrometric characterization as well.

**Figure 2.**
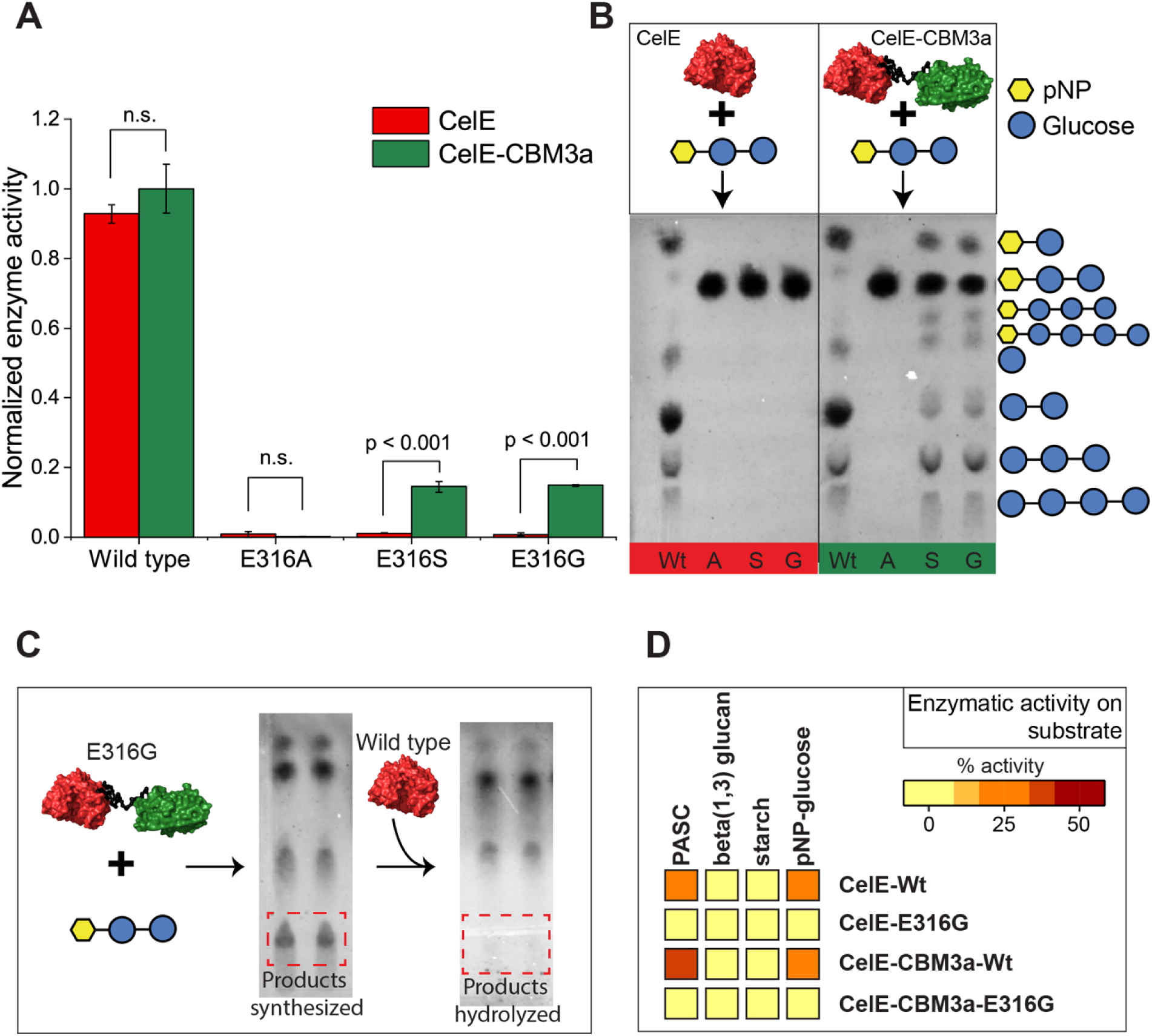
Carbohydrate binding modules (CBMs) aid in recovery of transglycosylation activity of nucleophilic mutants belonging to a family 5 glycosyl hydrolase (CelE). Catalytic nucleophile mutants of CelE domain shows improved transglycosylation activity when tethered to a family 3a carbohydrate binding domain (CBM3a). (A) The relative enzyme activities were estimated by measuring the release of pNP from pNP-Cellobiose (pNPC) as starting substrate. Here, 500 picomoles of each purified protein was incubated along with 2 µmoles of pNPC and the reaction was run for 4.5 h at 60°C reaction temperature. All reactions were run in triplicates with error bars representing 1σ from reported mean value. Mutation of catalytic nucleophile (E316) abrogated the activity of only the CelE catalytic domain (red color), while the E316G and E316S mutants of CBM3a tagged CelE showed ~20% pNP-release activity. (B) TLC image confirms the formation of longer chain oligosaccharides in the reaction mixture and the improved transglycosylation to hydrolysis product ratio (TG/HT) observed for the mutants. (C) The products formed by the mutant enzymes were completely hydrolyzed when supplemented with wild type CelE enzyme indicating that the glycosidic linkage between the products is β-retaining. (D) Activity assays of various enzyme on substrates with different linkages (e.g., β(1,4), β(1,3) and α(1,4)) further suggested that the transglycosylation reaction products contained β(1,4) linkages.

Clearly, the CBM3a domain fused to the CelE-E316S/G nucleophile mutants significantly (p<0.001, based on pNP release data shown in Figure 2A) improved hydrolytic and/or transglycosylation activity of the inactive catalytic domains towards pNPC substrate, compared to CelE-E316S/G domains alone, that ultimately led to the formation of cellotriose and cellotetraose along with minor amounts of cellobiose. Addition of exogenously added CBM3a to CelE-E316G did not show recovery of any transglycosylation activity (**SI Figure S3**), which suggested the subtly-timed and highly dynamic intra-domain interactions for the full-length chimeric E316G nucleophile mutants of CelE-CBM3a is likely very critical to recover transglycosylation activity. Also, increasing the added enzyme concentration over >2 orders of magnitude did not have a significant effect on the pNP-release activity of CelE-E316G unlike the CBM3a tethered CelE-E316G construct (**SI Figure S4**). This suggested that the pNP release rate for CelE-E316G was not limited by the relative enzyme to substrate concentrations in our assay conditions. Furthermore, the E316G nucleophile mutants of CelE-CBM3a gave no hydrolytic or transglycosylation activity on pNP-glucose alone as starting substrate suggesting that the CBM3a domain was somehow activating the ‘inactive’ CelE-E316G catalytic domain only in presence of pNPC based activated substrate (**SI Figure S5**). This very unusual behavior in β-(1,4)-glucan synthase activity could be clearly attributed to the presence of the tethered CBM3a domain that could have either potentially increased the local concentration of substrate around the CelE domain or could have facilitated inter-domain interactions between CBM3a-CelE to somehow play an active role in either stabilizing the substrate and/or product in the catalytic turnover step. For both these scenarios, we hypothesized that there were likely additional CBM3a residues participating in the catalytic turnover of the E316G mutant catalytic domain to result in β-retaining gluco-oligosaccharide products formed from a starting β-substrate (i.e., pNPC).

### Improved glucosynthase activity unique to CBM3a amongst other Type-A/B family CBMs

The peculiar glucan synthase activity results could be associated with increased local substrate concentration due to interactions of the carbohydrate-binding motifs of the CBM with the pNPC substrate, via sugar-π, π–π and hydrogen bonding interactions (Boraston et al., 2004; Boraston, 2005), that drive up the reaction rate of CelE-E316G domain. This was unlikely the reason responsible for the observed synthase activity since the reaction mixtures were homogenously well mixed and also the pNPC substrate has high water solubility (~50 g/L). Nevertheless, we hypothesized that if the increased substrate concentration is driven by the binding of pNPC towards CBM3a, this behavior could be possibly extended to other CBMs with similar affinity towards pNPC or cellobiosyl-like substrates (e.g., cellulose). Representative Type-A (CBM1 from *Trichoderma reesei*) and Type-B (CBM17 from *Clostridium cellulovorans*) CBMs were fused instead of CBM3a to CelE-E316G and tested for pNP release activity towards pNPC. Crystal structures of the three CBMs studied here (CBM1 PDB ID: 1CBH; CBM17 PDB ID: 1J84; CBM3a PDB ID: 1NBC) are shown in **SI Figure S6**. The reactions were run for 45 hours total (Figure 3) and the total pNP release was continuously monitored and is shown plotted in Figure 3A for multiple time points for all tested enzyme constructs. The wild type CelE-CBM constructs (i.e., with no E316G mutation) reached equilibrium within 2 hours of reaction at 60°C with comparable measured specific activities based on overall pNP release rate (Figure 3B). While the presence of CBMs did not largely impact the activity of CelE-WT domain, it was observed that only the CBM3a domain contributed to a ~60-fold increase (molar basis based on overall pNP release rate) in the specific activity of CelE-E316G catalytic domain. The CBM1 or CBM17 caused only a marginal 2 to 3-fold improvement in specific activity compared to the CelE-E316G catalytic domain alone.

**Figure 3.**
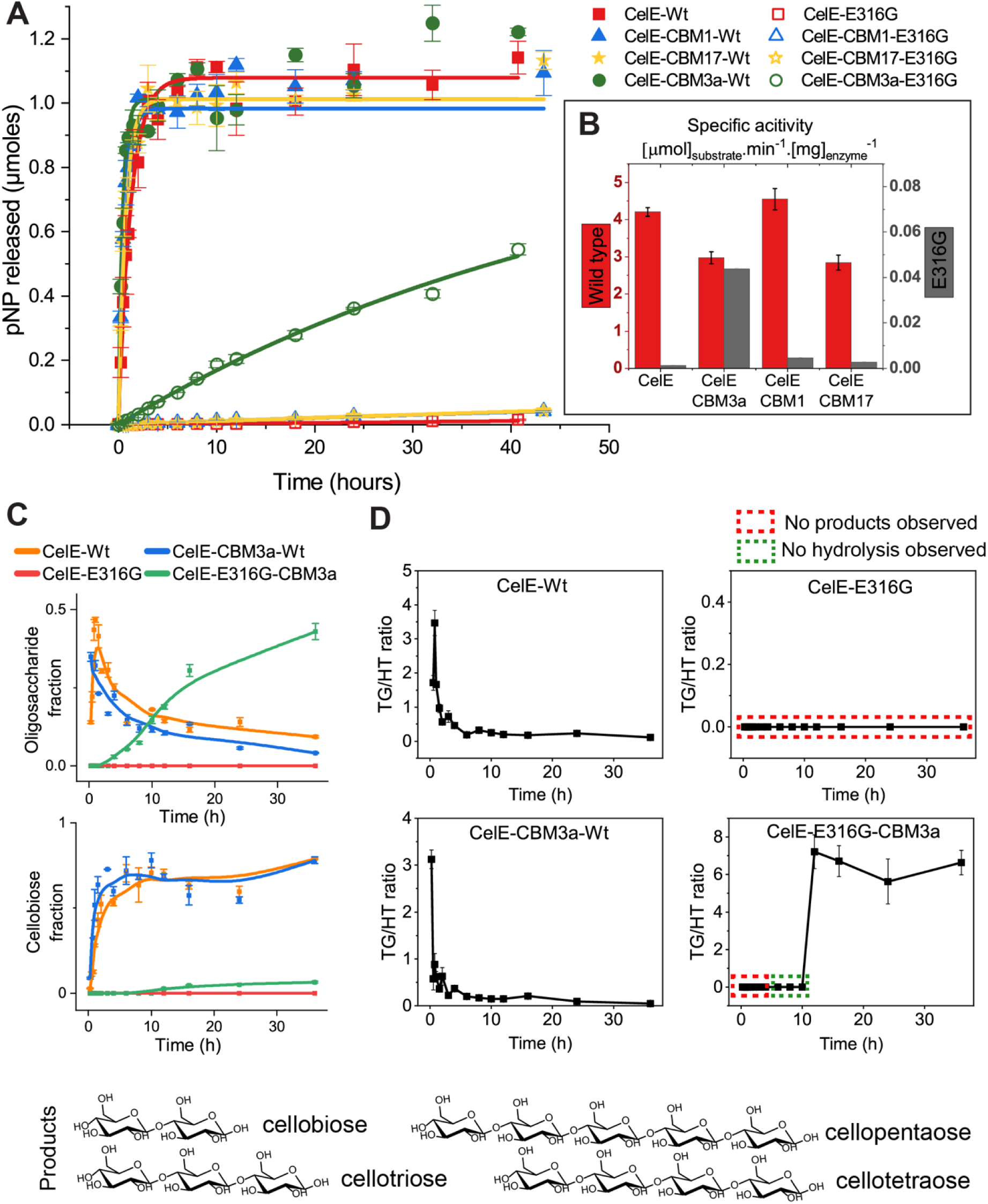
CBM3a enhances transglycosylation reaction rate of CelE nucleophile mutant by nearly two orders of magnitude unlike other CBM families. (A) CBM1 (Type A) and CBM17 (Type B), which have similar binding affinity towards cellobiose, were shown to not significantly impact the pNP-release activity of E316G mutant of CelE. End-point kinetic assays performed by measuring the pNP released over time showed an increased activity was seen only for the CBM3a tagged E316G CelE mutant. (B) The specific activities of all enzymes were estimated using the kinetic data for initial activity time points to show that there was a nearly 60-fold increase in the specific activity of CBM3a tagged E316G mutant compared to the CelE-E316G control. (C) Fractional concentration profiles for oligosaccharides and cellobiose formation are shown here based on product profiles measured for various timepoints. The most abundant product for wild type enzymes was cellobiose while for CelE-E316G-CBM3a construct the transglycosylation products are mostly dominant. (D) The transglycosylation/hydrolysis product (TG/HT) ratio for the four constructs indicate that the CelE-E316G-CBM3a is a highly efficient transglycosidase with a ~140-fold higher TG/HT ratio as compared to wild type counterpart as the reaction proceeds to equilibrium. During the initial rate, there were no hydrolyzed products observed and only oligosaccharides based products were seen. For these time points the TG/HT were not computed (indicated in green box). CelE-E316G mutant is not active and shows no transglycosylation or hydrolysis products and hence the TG/HT ratio was not computed (red box). Error bar indicates one standard deviation from reported mean values from biological replicates as highlighted in methods section.

Furthermore, in addition to monitoring pNP release, we also quantitatively characterized the total product profiles of transglycosylation and hydrolysis products for several enzyme constructs as shown in Figure 3C and **SI Figure S7-S8**. Interestingly, while CelE-E316G nucleophile mutant showed no significant change in reaction product profile over the entire duration, wild-type CelE-CBM3a showed formation of some transglycosylation products within the first 15-30 mins of the reaction that mostly disappeared as the reaction tended towards equilibrium giving rise to predominantly cellobiose as final hydrolysis product. On the other hand, CelE-E316G-CBM3a clearly showed a significantly higher concentration of multiple cellodextrins (along with major intermediate products like pNP-cellotriose and pNP-cellotetraose; see **Figure S8**) that increased in yield as the reaction slowly tended towards equilibrium. The fractional oligosaccharide product concentration profile for CelE-E316G-CBM3a shows that the majority of the products are indeed transglycosylation type products (~50%), while the remainder of the product fraction is unreacted pNP-cellobiose substrate along with only minor fractional concentration of cellobiose (<5%). This is unlike the product profile seen for wild-type enzymes that resulted in mostly cellobiose (>90%). Since measurement of various oligosaccharides requires the use of suitable chromatographic techniques (e.g., HPLC or TLC) which can prove to be tedious especially when dealing with a large number of mutants. Here, we note that total pNP release is directly correlated to the total oligosaccharide formation seen for the CelE-E316G-CBM3a construct. This allows easy high-throughput spectrophotometric based monitoring and rapid screening of mutants with improved transglycosylation prior to detailed TLC based analysis.

Figure 3D shows the total transglycosylation to hydrolysis (TG/HT) product ratio for the CelE-E316G-CBM3a construct and respective controls. We observe that wild type CelE constructs with or without CBM3a have similar TG/HT profiles. The initial products that are formed in the reaction using wild-type enzymes are transglycosylation products that nearly instantly get hydrolyzed into cellobiose as the reaction proceeds to equilibrium. This can be clearly seen as the TG/HT ratio decreases rapidly to less than 0.2 within few hours of reaction. No products were seen for CelE-E316G mutant and hence the TG/HT ratio could not be computed for this construct. On the other hand, the presence of CBM3a tethered to CelE-E316G facilitated the transglycosylation activity of CD to form transglycosylation products initially (<10 hours) and virtually no hydrolysis products during this period. Marginal amounts of hydrolyzed products are subsequently observed as the reaction proceeds beyond 10 hrs. Ultimately, as the reaction proceeded to equilibrium, the CelE-E316G-CBM3a construct gave TG/HT product ratio ranging between 40 to 140-fold higher than the values obtained for the CelE-CBM3a-Wt control.

Lastly, although no direct measurements to characterize binding affinity of pNPC or cellobiose to CBMs was conducted here, the crystalline cellulose binding affinities for both CBM3a and CBM1 are very similar and in the µM affinity range (Georgelis et al., 2012; Linder and Teeri, 1996). While, CBM17 has been shown to bind specifically to both soluble sugars and insoluble amorphous cellulose, albeit with much weaker affinities in the mM-to-µM range (Boraston et al., 2000; Georgelis et al., 2012). Clear differences observed in the activities of CelE-E316G fused to different CBMs with similar binding characteristics suggest that the observed β-glucan synthase activity seen was not necessarily driven primarily by increased local substrate concentration. Instead, there are likely additional subtle and highly specific interactions between the CBM3a and CelE domains that are critical to recovering catalytic activity for the CelE-E316G nucleophile mutant. Closer inspection of the available crystal structures of the three CBMs studied here (CBM1 PDB ID: 1CBH; CBM17 PDB ID: 1J84; CBM3a PDB ID: 1NBC) revealed the presence of an additional hydrophobic cleft (**SI Figure S6**) in CBM3a which might assist in substrate and/or product stability in synergy with the CelE active site as scaffold for synthase activity. The original 42aa linker between the CelE and CBM3a is moderately flexible with multiple threonine residues that likely facilitates interactions between the two domains. Additionally, since the linker is native to CBM3a from a cellulosomal bacterial system, it is likely that the linker dynamically folds to form compact structures facilitated by CBM3a specific-interactions not possible with other CBMs (i.e., CBM1, CBM17). This could also explain why CBM3a alone gave the highest measured β-glucan synthase activity over CBM1 and CBM17.

### Specific inter-domain interactions of CelE-Linker-CBM3a facilitates transglycosylation

We hypothesized that specific inter-domain interactions between CelE-E316G and CBM3a due to the flexible native 42aa linker likely facilitated transglycosylation activity for the chimeric E316G nucleophile mutants of CelE-CBM3a. The compactness/flexibility of the domains and conformer ensemble populations could be predicted using small angle x-ray scattering (SAXS) experiments in the presence and absence of substrate (Figure 4). SAXS measurements were conducted for CelE-CBM3a and CelE-E316G-CBM3a in the presence and absence of pNPC substrate. The SAXS profiles of CelE-E316G-CBM3a, after solvent background subtraction, are shown as double log plot in Figure 4A and the p(r) distribution plots are illustrated in Figure 4B. The scattering intensity for the two domain CelE-E316G-CBM3a protein alone appears to have a sharp drop at the mid-q to high-q range, while the same protein samples quenched with excess pNPC substrate showed a more gradual drop. These results indicate that the ensemble is composed of a limited set of conformations for the protein only samples, however, the samples which have both the protein-substrate display a large number of dynamic conformations. The asymptotic behavior of SAXS intensity decay in the Porod regime following the Guinier region provides structural information about the shape of the scattering sample.

**Figure 4.**
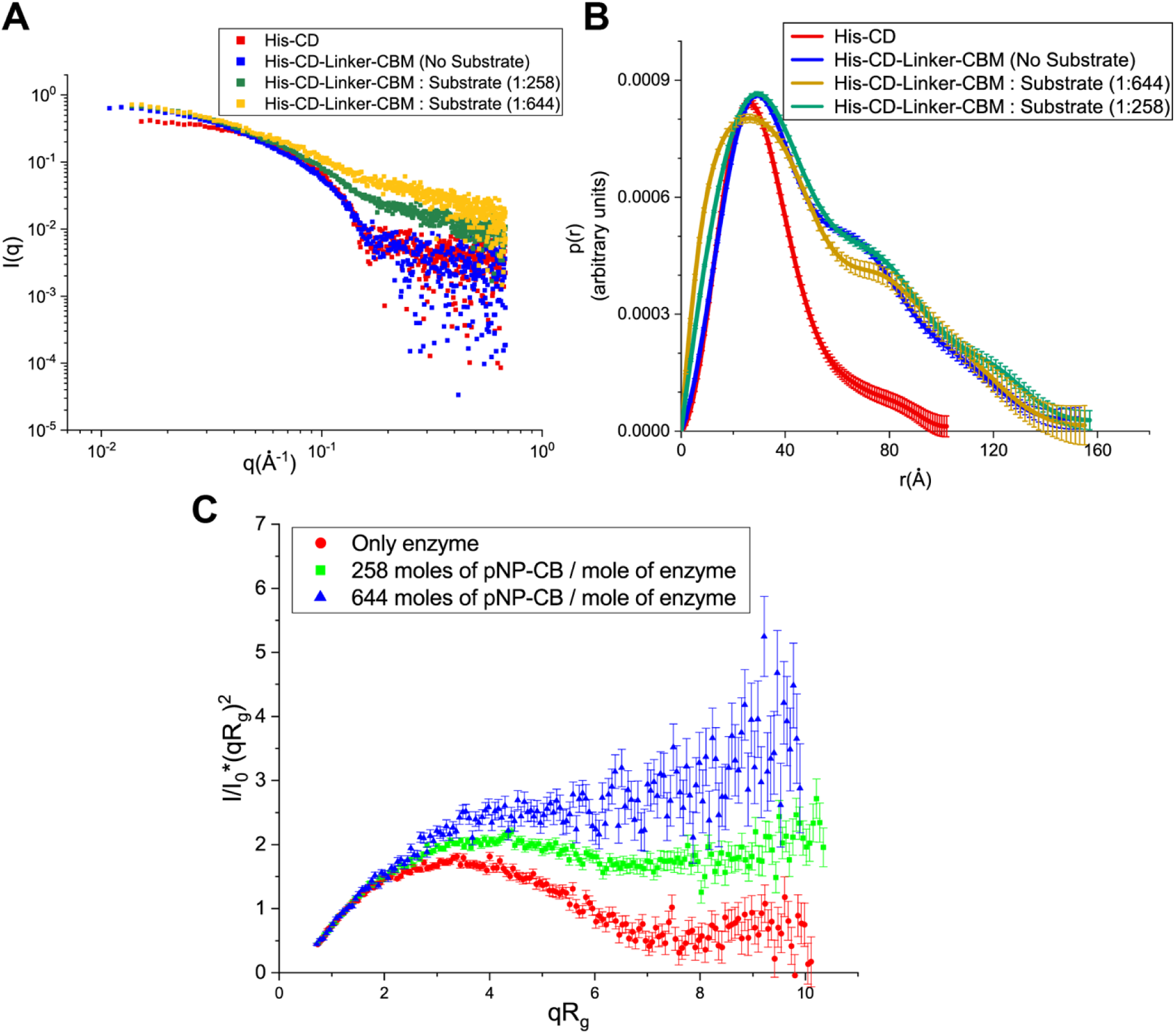
Small angle X-ray scattering indicates the presence of dynamic flexibility between the CBM3a and CelE-E316G domains. (A) Original SAXS profiles and (B) analyzed p(r) distribution profiles of CelE-E316G and CelE-E316G-CBM3a enzyme with and without pNP-cellobiose are shown here. The distance distribution function (p(r) curves) for each sample were generated from original SAXS data to estimate the real space radius of gyration (Rg) and maximum dimension (Dmax). (C) Kratky plots for CelE-CBM3a-E316G mutant enzyme with and without pNP-cellobiose (pNP-CB) highlights the presence of varying degrees of flexibility in the presence of pNP-CB substrate.

The Kratky plot (i.e., q^2^R_g_I(q)/I_0_ vs. qR_g_ plot) can qualitatively assess the compactness and degree of unfolding of multidomain proteins. The Kratky plots for two domain proteins shown in Figure 4C demonstrate that CelE-E316G-CBM3a exists as a compact structure in solution in the absence of any substrate (and see **SI Table S2** and Figure 4B for additional details on SAXS analysis of other controls/samples). Upon addition of the pNPC substrate, the inter-domain flexibility or ‘breathing’ motion between two domains increases compared to no substrate samples, indicative of a multi-domain protein with a highly flexible linker domain. Furthermore, the dynamic motion between the two domains increases as the concentration of pNPC substrate added relative to protein is increased. This suggests that either; (i) initially a more compact CelE-E316G-CBM3a opens up to make the active site more accessible and the flexibility between the two domains enable the domains to come closer to stabilize substrate interactions and facilitate synthase activity, and/or (ii) the energy released during the slow transglycosylation reaction taking place causes the dynamic flexibility between two domains. In either case, a subset of ensemble of protein conformations likely conform close proximity of the two domains. An ensemble of conformations taken by multi-domain protein could be readily generated using the ensemble optimization method (EOM). EOM analysis of the experimental SAXS profiles was used to predict the model shapes and the fraction of conformations that exhibit close proximity of CelE and CBM3a was determined. Multiple conformations were observed where the CBM3a domain was very close to the native CelE active site cleft. The presence of a long flexible linker chain (42 aa) likely facilitated the dynamic interaction of CBM3a close to the CelE substrate binding site cleft. Furthermore, the native linker was found to be critical for facilitating the inter-domain interactions driving the glucan synthase activity (Figure 5). This was evidently seen from the results shown in Figure 5A where the 42 aa linker was truncated to 6 aa, 11 aa, and 21 aa. The shortest linker (6 aa) resulted in a significant decrease in the relative pNP-release activity, while the 21 aa linker showed higher activity compared to even the original native 42aa linker. An optimal linker length might help stabilize the inter-domain interactions by reducing the number of extended conformations and leading to more compact structures. In the control experiments for wild-type CelE and CelE-CBM3a constructs with no nucleophile site mutations, the impact of the linker length was insignificant on the wild type enzyme pNPC hydrolytic activity indicating that the linker truncations did not likely affect protein folding. Along with the linker length analysis, the linker flexibility was also modified by altering the 42aa linker sequence composition to make the linker either highly flexible or highly rigid compared to the native linker for the same linker length (e.g., more flexible (F2) > F1 >> Native Linker >> R1 > (R2) more rigid). The relative activity profiles of these highly flexible and rigid proteins are shown in Figure 5B, which confirms that the extreme ranges of linker flexibility deleteriously impact the CBM-CD domains from forming a productive complex necessary for achieving optimal synthase activity. Though, it seems clear that the more flexible linkers (F1 and F2) give marginally higher pNP-release activity activity than the more rigid linkers (R1 and R2).

**Figure 5.**
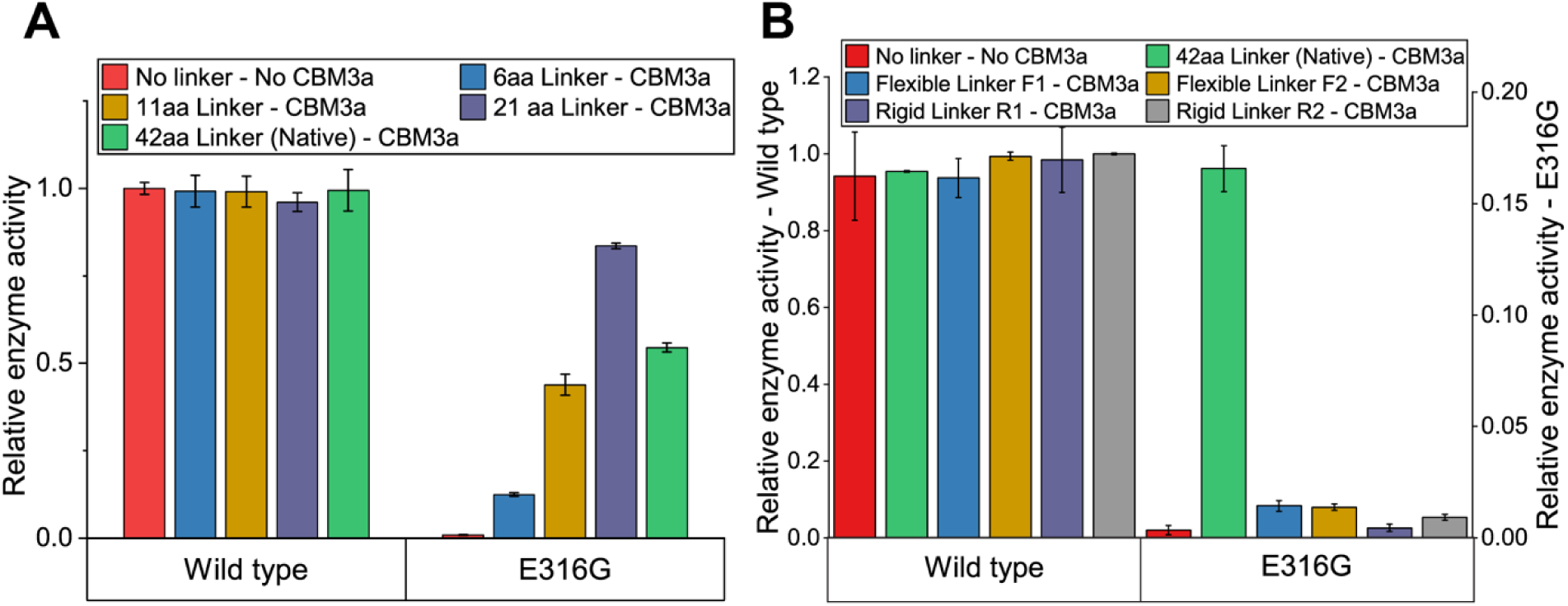
Inter-domain interactions of CelE and CBM3a domains critical for observed transglycosylation activity is facilitated by optimum linker size and sequence structure. (A) Biochemical assays to verify the critical role of inter-domain interactions facilitating transglycosylation activity were performed by varying the size of the linker between the CelE and CBM3a domains. A very short linker (6aa linker) showed reduced activity as compared to the original 42aa native linker. Furthermore, there appeared to be an optimal linker length (21aa) that strengthened the interactions between the two domains to further improve transglycosylation activity compared to the native 42aa linker. (B) Flexibility of the 42aa long linker was varied to become either highly flexible (F1 and F2) or highly rigid (R1 and R2) and the corresponding transglycosylation activities of the resultant mutants are highlighted here. The native linker of CBM3a plays a very important role in aiding the inter-domain interactions unlike other similar sized linkers of varying sequence structure. Nevertheless, flexible linkers gave always higher activities than more rigid linkers.

### Both CelE and CBM3a substrate binding cleft residues are critical for efficient transglycosylation

Formation of typical transglycosylation product intermediates, though mostly short-lived once the reaction reaches equilibrium, for wild-type β-retaining GHs to form β-retaining products like pNP-cellotetraose or cellotetraose necessitates the formation of a covalent GEI (**SI Figure S9**). However, in our study all three nucleophilic site mutants (i.e., E316G/E316A/E316S) of CelE were inactive in the absence of a tethered CBM3a (Figure 2B). We speculated that the hydrophobic pocket on the underside of CBM3a could also facilitate in the departure of the pNP leaving group sandwiched between the pNPC substrates bound to the CelE-E316G native glucan substrate binding cleft. We suspected that CelE-E316G provided a suitable substrate binding scaffold that likely oriented the donor and acceptor glycone groups adjacent to each other to facilitate intermolecular glycosyl transfer via either a poorly understood S_N_i-like mechanism or S_N_2-like mechanism due to the presence of a currently unknown pseudo-nucleophile residue. Either ways, in light of all experimental results discussed earlier, it was only possible for a β-retaining transglycosylation product like pNP-cellotetraose to be formed if the CelE-E316G-CBM3a domains worked together to employ a suitable catalytic glycosyl transfer step.

The CelE-E316G-CBM3a protein model based on a structure predicted using SAXS and the available crystal structures CelE (PDB ID: 4IM4) and CBM3a (PDB ID: 1NBC) was therefore analyzed further to explore the role of key CD/CBM residues on the observed transglycosylation reaction (Figure 6). The additional hydrophobic cleft on the underside of CBM3a could potentially be involved in facilitating the glycosynthase reaction through a poorly understood reaction mechanism. However, while there was no similar CBM3a cleft structural analogue on the other Type-A/B CBMs we investigated, there was a minimal but significant ~2-fold improvement in pNP release activity observed for the other two constructs (i.e., CelE-E316G-CBM1 and CelE-E316G-CBM17) compared to CelE-E316G alone. This suggests that stabilization of the pNP leaving group via the cellulose-binding planar surfaces of each CBM (see **SI Figure S6** and Figure 6B), including CBM3a, is also likely feasible. But this alternative mechanism of pNP leaving group stabilization would be likely less favorable, in keeping with the relatively very slow rate of reaction observed for CBM1/CBM17 tethered CelE-E316G. For these reasons, further targets for mutational studies aimed at disrupting the residues in the CBM3a hydrophobic pocket, and in particular the aromatic residues known to be often involved in sugar-aromatic stacking interactions. In addition, the aromatic residues within the CD substrate binding cleft (Y270, Y273, W203, W349, H268) and active acid/base residue (E193) that could potentially participate in the reaction were also selected (Figure 6A). All identified residues (five on CBM3a and six on CelE domains) were mutated to alanine, creating double mutant constructs (along with CelE-E316G included in each case). Similarly, twelve individual single mutations were introduced while keeping the catalytic nucleophile of CelE intact. These additional twelve controls were used to account for any potential activity loss due to protein misfolding. While all the control samples (with single point mutations on CBM3a alone) showed identical activity as the wild type CelE-CBM3a enzyme (Figure 6C), the double mutations introduced resulted in a significant loss in pNP-release activity compared to CelE-E316G-CBM3a (Figure 6D). Mutation of each CBM3a residue decreased the pNP release activity by at least 50% when compared to CelE-E316G-CBM3a, which was also directly correlated to the higher molecular weight transglycosylation products formed. All of the aforementioned residues of CBM3a together could contribute to stable docking of pNPC substrate or pNP leaving group along with the CelE substrate binding site scaffold to a certain extent and no mutation alone caused a complete loss of transglycosylation activity.

**Figure 6.**
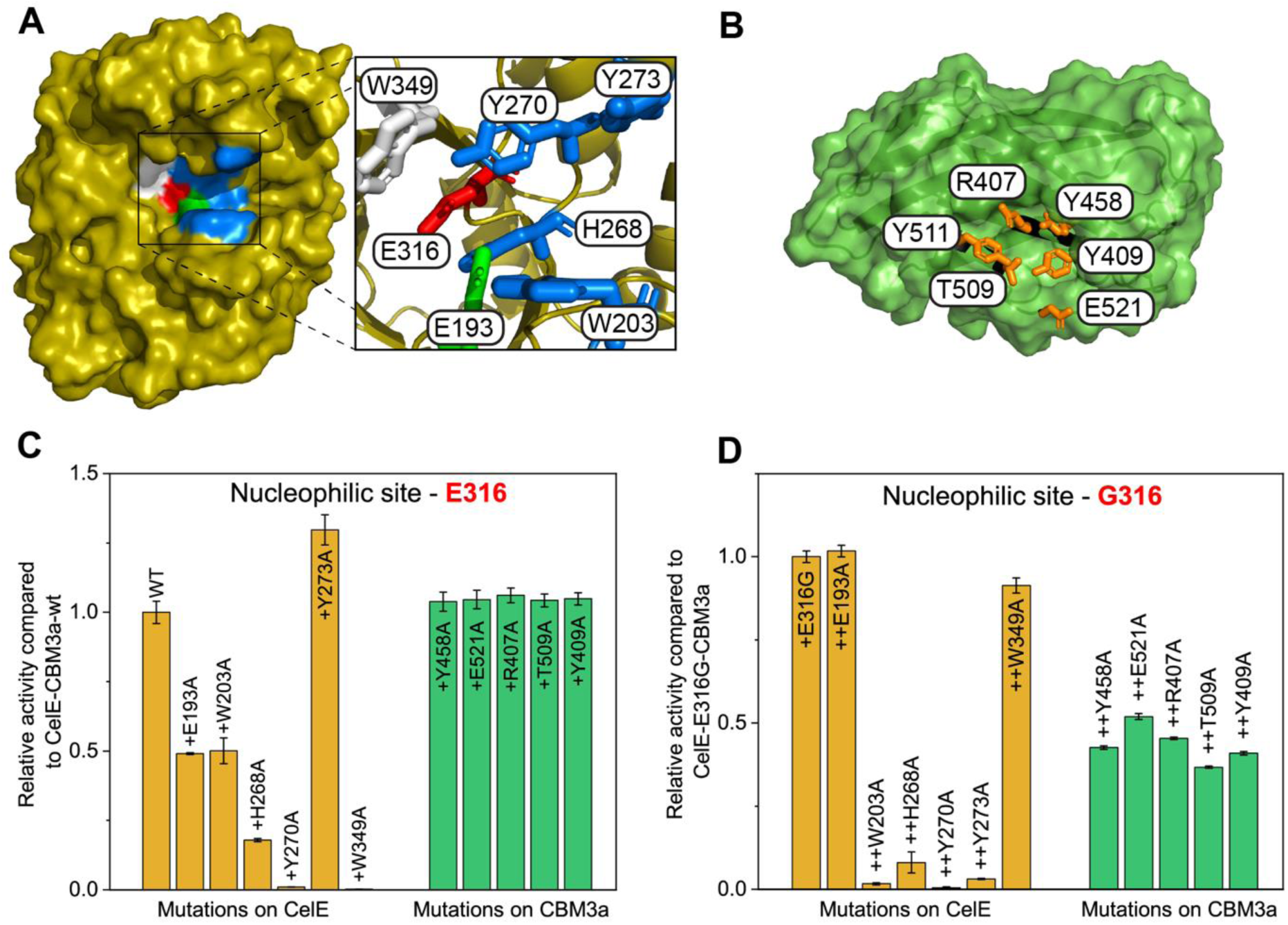
Impact of critical amino acid residues identified within the hypothesized CelE-E316G-CBM3a enzyme-substrate complex that impact transglycosylation activity. (A) Active site cleft of native CelE enzyme is shown here. The catalytic nucleophile (E316 – red), catalytic acid/base (E193 – green), negative subsite (W349 – white) and positive subsites (Y270, Y273, W203 and H268 – blue) are highlighted. (B) Key residues within the hydrophobic cleft of CBM3a are illustrated here (R407, Y458, Y409, E521, and T509 – orange). (C) Relative activity of single alanine point mutations of the key identified residues of CelE and CBM3a with native nucleophile as compared to CelE-CBM3a-wt is present here as bar graphs. (D) Activity of the double mutants containing alanine mutation of key residues with nucleophile also mutated to glycine (E316G) are shown as relative activity compared to CelE-E31G-CBM3a.

MAFFT sequence alignment of the CelE protein sequence and CBM3a protein sequence to protein sequences of their respective GH/CBM families showed that these interacting residues are highly conserved throughout these families (**SI Figure S10**), and we propose that similar mechanisms are likely to operate in similar GH-CBM chimeric constructs. Preliminary experimental studies have indeed confirmed that fusion of the CBM3a-42aa linker domain to nearly a dozen other phylogenetically related GH5 families (and all with corresponding nucleophile site mutations to glycine) resulted in active transglycosidases as seen here with CelE-E316G-CBM3a. Representative results for one other homologous GH5 family protein fused to CBM3a are shown in **SI Figure S11**. Furthermore, similar detailed time-course assays (**SI Figure S8** and Figure 3C) and future kinetic modeling of the reactions hypothesized to form the complex products suite for CelE-E316G-CBM3a system for other structurally homologous GH-CBM constructs will help identify the rate-limiting steps relevant to the targeted formation of other desired intermediate products (e.g., pNP-cellotetraose).

Our current results also suggest the possibility of either an S_N_i or even a S_N_2 type glycosyl transfer mechanism that is possible for the current chimeric constructs. The S_N_i-like mechanism is a substrate assisted mechanism where the protein does not actively form any covalent bond with the substrate and only aids in creating bond strains through non-covalent binding interactions between the protein active-site scaffold and the substrates. Since all retaining enzymes operating with an S_N_2 type mechanism function with the help of two catalytic residues; the nucleophile and acid/base residues, we performed additional mutational studies to study the impact of the acid-/base residue (Figure 6D). As expected, mutating the acid/base residue (E193) to alanine abrogated the pNP-release hydrolytic activity of CelE-E193A by ~50% (compared to control enzyme construct with an intact catalytic nucleophile or CelE-CBM3a) confirming the likely participation of this predicted acid/base residue in the retaining mechanism of native CelE. However, introducing the E193A mutation into CelE-E316G-CBM3a construct did not cause any change in pNP-release activity suggesting that the traditional acid/base residue does not likely participate in the catalytic reaction for our chimeric glucan synthase mutants. Similar results regarding the non-participatory role of traditional acid/base residues on another S_N_i-like mechanism have been recently reported as well (Iglesias-Fernández et al., 2017). Interestingly, single point mutation of several other active site residues of CelE alone (Figure 6C), which were in close vicinity to the native GEI transition state complex, further reduced the pNP-release activity by 90% or even higher (e.g., Y270A, W349). These supplementary results further suggests the possibility of other pseudonucleophile residues like Y270 that could facilitate transglycosylation/hydrolysis activity via a classical S_N_2 mechanism or even a transition state stabilizing role for a S_N_i-like mechanism as well, as shown recently for another GH family 1 glycosynthase (Iglesias-Fernández et al., 2017). While, clearly all these residues play a key role in facilitating transglycosidase activity facilitated by the CBM3a domain, the actual reaction mechanism of glycosyl transfer will be unraveled in the near future via ongoing detailed protein mutagenesis and molecular simulation studies.

## Discussion

GHs and GTs are the two major classes of enzymes that are known to catalyze the hydrolysis and/or synthesis of glycosidic linkages between carbohydrate moieties. Although both these enzyme classes are functionally different, their mode of action on glycosidic bonds follows similar basic design principles (i.e., donor glycone with a good leaving group is ‘activated’ in the enzyme active site to generate a short-lived intermediate or stable GEI and next a suitable acceptor molecule is attached to the activated/intermediate donor glycone group). This could explain why gene annotation/classification of these enzymes based on sequence similarities and protein folds often results in significant overlap between the two enzyme classes (Hidaka et al., 2004; Lombard et al., 2014; Roston et al., 2014). However, few studies have explored the role of CBMs on transglycosylation activity of GHs and none have so far explored the possibility of the reaction mechanism directly employing CBMs (Codera et al., 2015; Mizutani et al., 2012; Stockinger et al., 2015). While CBMs or lectin-like domains have been never reported previously to directly participate in the catalytic reaction step for glycosidic bonds synthesis by GHs, these domains have been shown to play important roles in the functioning of several GTs. GTs are often associated with one or multiple CBMs/lectins-like domains on either the N- or C-terminal ends, where these domains participate in substrate recognition and in some cases participate in active site cleft formation to facilitate biocatalysis. For example, GTs responsible for mucin biosynthesis, namely N-acetyl galactosamine transferases (GalNAc-T1 (Fritz et al., 2004) and GalNAc-T2 (Lira-Navarrete et al., 2015)), belonging to the GT27 family have a CBM13 domain tethered to the C-terminus of the catalytic domain. The two domains were shown to dynamically interact with each other forming an active site cleft to facilitate addition of a N-acetyl-galactosamine donor group to an acceptor polypeptide chain (Lira-Navarrete et al., 2015). The linkers in multi-domain GTs have also co-evolved with CBMs/lectin-like domains to stabilize these ’active’ inter-domain conformations that enable efficient glycosidic bonds synthesis. But, we currently have a poor understanding of the dynamic interplay between multiple domains due to the inherent lack of available crystallographic data or the subtle impact of such dynamic domain interactions on overall catalytic turnover rates. The linker regions between most GHs and CBMs also tend to be much more flexible than GTs to allow for adjustments to complex structural motifs often found in polysaccharide and oligosaccharide type substrates (e.g., in plant cell walls) (Currie et al., 2013; Receveur et al., 2002). Nevertheless, multi-domain interactions are widely extant in CAZymes, and therefore likely play a significant but poorly understood role in modulating both GTs synthase and GHs hydrolase (and transglycosidase) activity (Albesa-Jové and Guerin, 2016). Better understanding how similar GH-CBM chimeras allow subtle fine-tuning of transglycosylation mechanism that allow selective formation of glyco-synthase versus glyco-hydrolase based reaction products could help improve cellulosic biofuel production using engineered CBP microbes. Considering that most cellulolytic bacterial CAZymes are often appended to CBMs (Brumm, 2013), it is likely that engineered cellulosomal enzyme complexes could utilize similar mechanisms to generate cello-oligosaccharides for synergistic uptake by CBP bacteria (like *C. thermocellum*) could allow for more efficient fermentation of cellulosic biomass into ethanol (Lu et al., 2006).

Here, we reported novel chimeric CBM-based enzyme designs that provide an unexplored route for chemoenzymatic synthesis of bespoke glycans like cellodextrins and pNP-based oligosaccharides. We have found that, similar to multi-domain GTs, engineered chimeric GH 5 catalytic domain scaffolds (in the absence of any true nucleophile residue) tethered by a linker to CBM-like domains can dynamically interact to form ‘active’ GH-CBM complexes that facilitated synthesis of glucose based oligosaccharides from activated donor sugar monomers. We observed that the total transglycosylation to hydrolysis (TG/HT) product ratio for the CelE-E316G-CBM3a construct was 40 to 140-fold higher than the values obtained for the control wild-type enzymes. Clearly our chimeric mutant is a highly efficient transglycosidase based on TG/HT metrics established for other CAZyme systems (Lundemo et al., 2013; Nordvang et al., 2016). Previous studies on another glucosidase have reported similar order of magnitude (70-fold) increase in TG/HT ratio but only after extensive mutagenesis and directed evolution of the wild-type enzymes (Kone et al., 2008), which further highlights the advantages offered by our current approach. This approach could be used to also target highly efficient synthesis of other non-cellulosic glycan polymers using readily accessible pNP-glycosides or naturally available aromatic leaving group based glycosides as substrates (Gantt et al., 2013). However, future work will need to also focus on fully unraveling the reaction mechanism and showcasing how similar CBM-/lectin-like domain assisted mechanism could be prevalent in other GH families as well.

In the case of CelE-E316G-CBM3a construct, the CBM3a domain is speculated to stabilize the leaving group (pNP) near the active site substrate binding cleft of CelE allowing for the first transglycosylation reaction to take place forming pNP-cellotetraose or similar pNP based oligosaccharides. While, similar transglycosylation products are likely also formed for native CelE-CBM3a, these products are nearly instantly degraded to produce the equilibrium dominant products (i.e., pNP and cellobiose). In the absence of the native nucleophilic residue (E316), significantly higher concentration of transglycosylation products are seen even as the reaction tends towards equilibrium. Here, the rate-limiting step is hypothesized to be the formation of the active CelE-E316G-CBM complex that facilitates the synthesis of the higher molecular weight transglycosylation products. This hypothesis is consistent with the critical importance of the linker identified that likely facilitates highly concerted dynamic, but poorly understood, interactions between the GH, CBM, and substrate. Since S_N_i-like/S_N_1 and S_N_2 type mechanisms are often seen to operate in retaining and inverting GTs (Lairson et al., 2008), respectively, it is likely that similar mechanisms could operate in GHs as well. However, only S_N_1 and S_N_2 type mechanisms have been reported for GHs, but with one recent exception (Iglesias-Fernández et al., 2017). However, it is also possible that there is a likely pseudo-nucleophile residue that participates in a conventional S_N_2-like reaction mechanism (i.e., after mutation of E316G). Nevertheless, our findings suggest that researchers should more closely investigate the role of ‘true’ nucleophilic residues for each specific GH families, which is likely substrate-dependent as well.

## Acknowledgments

SPSC acknowledges support from the US National Science Foundation CBET Division (Award No. 1704679), ORAU 2016 Ralph E. Powe Award, ORNL Neutron Sciences User Facility, and Rutgers School of Engineering. SVP acknowledges the support of DOE Office of Science, Office of Biological and Environmental Research resource, the Center for Structural Molecular Biology (CSMB) and Office of Basic Energy Sciences, Scientific User Facilities for the BioSAXS 2000 resource, operated by the Oak Ridge National Laboratory. We are very grateful to other members from the Chundawat research group for their critical feedback and contributions during the course of this project. We thank Madhura Kasture for her contributions towards generating the activity data for homologous proteins. Lastly, the authors thank Dr. Heather Mayes and Tucker Burgin at the University of Michigan for helpful discussions regarding the S_N_i/S_N_2 reaction mechanism. Conflict of interest note: CKB and SPSC have filed a US provisional patent application (No. 62/719,963 filed by Rutgers University on Aug 20th 2018).

This manuscript has been authored by UT-Battelle, LLC, under Contract No. DE-AC05-00OR22725 with the U.S. Department of Energy. The United States Government retains and the publisher, by accepting the article for publication, acknowledges that the United States Government retains a non-exclusive, paid-up, irrevocable, world-wide license to publish or reproduce the published form of this manuscript, or allow others to do so, for United States Government purposes. The Department of Energy will provide public access to these results of federally sponsored research in accordance with the DOE Public Access Plan (http://energy.gov/downloads/doe-public-access-plan).

## Supplementary Online Material

Supporting Information (SI) appendix pdf is provided online with supplementary methods/text and supplementary results (as SI Figures S1-S11, Tables S1-S2).

