## Supplementary Information for "Carbohydrate binding domains facilitate efficient oligosaccharides synthesis by enhancing mutant catalytic domain transglycosylation activity"

**This PDF file includes:**

Supplementary Text (including Materials and Methods)
Figures S1 to S11
Table S1 to S2
SI References

### Supplementary Text

### Materials and Methods

***Bacterial strains and plasmids:*** All the chemicals, reagents and solvents were purchased from Fisher Scientific and Sigma-Aldrich and used without purification. *E. coli* strains used for cloning and protein expression were E. cloni® 10G (Lucigen, WI, USA) and BL21-CodonPlus-RIPL [λDE3] (Stratagene, Santa Clara, CA, USA) respectively. The plasmids, pEC-CelE, pEC-CelE-CBMs (CBM3a, CBM1 and CBM17) were kindly provided by the Fox lab at UW Madison. 2X Phusion high fidelity PCR master mix (0.04 U/µL Phusion DNA polymerase, 400 µM dNTPs, 2X Phusion HF buffer, 3 mM MgCl2) was purchased from Thermo Fisher Scientific (USA) and restriction enzymes were procured from New England BioLabs Inc. (USA). The primers used for site directed mutagenesis (SDM), sequence and ligation independent cloning (SLIC) and sequencing reactions were obtained from Integrated DNA Technologies, Inc (USA). Successfully cloned plasmids were isolated from E. cloni® 10g cells using IBI Scientific (USA) plasmid extraction kit and the sequence was confirmed through Sanger sequencing performed by Genscript Inc. (NJ, USA). The carbohydrate substrates used in enzyme assays were purchased from Carbosynth USA.

***Design and cloning of constructs***: The site directed mutagenesis of catalytic domain (CelE) and carbohydrate binding domain (CBM3a) was performed using Stratagene protocol to create point mutations. Polymerized chain reaction was performed using 1X Phusion master mix, template DNA and 0.5 µM of two complementary primers (**SI Table S1A**). After 20 cycles, the reaction mixtures were incubated with Dpn1 in 1X cut smart buffer at 37°C for 1 hr and transformed into E. cloni 10G competent cells and plated on LB agar plates supplemented with Kanamycin (50 µg/ml) to get selectively transformed colonies. Plasmids were isolated from individual transformed colonies and sequenced to confirm the nucleotide identity at the mutational site. Sequence and Ligation-Independent Cloning (SLIC) protocol was used to modify (insert or delete) the linker region between CelE and CBM3a domains (1). PCR amplifications of vector DNA and insert DNA were performed using their corresponding SLIC primers (**SI Table S1B**). The vector and insert PCR products after verification for amplification using DNA gel electrophoresis were washed to remove the unreacted nucleotides (dNTPs). The reaction mixture containing 0.025 pmol of vector and 0.0625 pmol insert purified products (2.5:1 insert: vector ratio) were digested using Dpn1 at 37°C for 1 hour. Dpn1 digested products were reacted with 1.5U of T4 DNA polymerase at 25°C for 5 mins and immediately placed on ice. Entire reaction mixture was transformed into E. cloni 10g cells and plated to get the colonies. The transformant colonies from the plate were screened using PCR amplification for identifying constructs with correct insert size observed in agarose gel electrophoresis. Plasmids from the selected colonies were isolated and sequenced to verify the nucleotide sequence. All the sequencing verified plasmids obtained from cloning were kept at -80°C and their corresponding transformed E. cloni cells were stored in 15% glycerol stocks at -80°C.

***Protein expression and purification:*** The successfully cloned and verified plasmids were transformed into *E. coli* BL21 cells for protein expression. Transformed colonies were inoculated into 25 ml of LB media supplemented with kanamycin (50 µg/ml) and grown at 37°C for 16 hrs. The 25 ml of overnight grown culture was transferred to 500 ml of fresh LB media with kanamycin and incubated at 37°C until mid-exponential phase (OD 0.4-0.6) was reached. At this point, protein expression was induced by adding 0.5 mM IPTG and protein induction was carried out at 25°C for 20 hrs - 24 hrs. The cells were harvested by centrifugation and the cell pellet was re-suspended in lysis buffer (20 mM sodium phosphate, 500 mM NaCl, 20% glycerol; pH 7.4) supplemented with lysozyme (10 µg/ml) and protease inhibitor cocktail (EDTA, E-64 and benzamidine). The cells were lysed by sonication at 4°C for 5 min with 10-s on-bursts and 30-s off periods. The cell debris was separated by centrifuging at 12000 g for 60 min at 4°C and the supernatant containing the protein of interest was collected and purified by immobilized metal affinity chromatography (IMAC) on Bio-rad NGC chromatography system using $Ni^{+2}$-NTA based column (GE Healthcare). The column was first equilibrated with binding buffer (IMAC A: 100 mM MOPS, pH 7.4, containing 10 mM imidazole and 100 mM NaCl) allowing the his-tag protein to be bound to the column when the cell lysate was passed through it. The column was washed with IMAC A along with an additional wash of 95% IMAC A and 5% elution buffer (IMAC B: 100 mM MOPS, pH 7.4, containing 500 mM imidazole and 100 mM NaCl) to remove the strongly bound non-specific proteins. The desired protein was eluted in 100% IMAC B buffer and buffer exchanged into 10 mM MES pH 6.5 using PD-10 desalting columns (GE Healthcare). The protein concentration was estimated by absorbance measurements at 280 nm in a spectrophotometer. The molecular weights and purity of the eluted proteins were confirmed by SDS-PAGE and matched to predicted translational products (**SI Figure S1 and Text S1**).

***End point activity assay***: The activity assays of CelE-CBM3a proteins on 4-nitrophenyl-β-D-cellobiose (pNP-cellobiose or pNP-CB) were performed in 200 µl reaction volume. 500 pmol of protein was reacted with 2 µmoles of pNP-CB in 50 mM MES pH 6.5 buffer at 60°C for 7 hours with shaking at 400 rpm. 10 µl of reaction mixture was taken at 3 hr (or 4 hr) and 7 hr and added to a microplate containing 100 µl of 0.1M NaOH and 90 µl of DI water. The absorbance of the microplate at

405 nm was measured and pNP released was estimated with the help of a calibration curve built for pNP standards and $\lambda_{405}$ absorbance.

***Kinetic activity assay***: Kinetic studies on the activity of enzyme on pNP-cellobiose were carried out at 60°C in MES buffer (50 mM, pH 6.5). The 150 µl reaction mixture consisted of 1.5 µmoles of pNP-CB and 100 pmoles of protein. Three replicates of individual reaction mixtures for every time point was setup. The reaction was quenched after each time point by denaturing the protein at 95°C for 5 minutes. The denatured tubes were centrifuged and 10 µl of supernatant was added to microplate well containing 100 µl of 0.1M NaOH and 90 µl of DI water to measure the pNP absorbance at 405 nm. Samples were collected for 8 proteins and 1 reaction blank for 17 time points spanning from 0 hr to 42 hrs. The unknown pNP absorbance was estimated using the calibration curve built using pNP standards and $\lambda_{405}$ absorbance.

***Quantitative Thin Layer Chromatography (TLC) analysis***: Reaction mixtures were analyzed by TLC using Analtech P21521 Silica Gel GHLF TLC plates for quantitative analysis of all hydrolysis and transglycosylation reaction products. The mobile phase used for TLC was ethyl acetate: 2-propanol: acetic acid: water (at 3:2:1:1 v/v ratios). Several standards were also run on the TLC plate to determine the unknown detected spots in reaction sample based on retention factor ($R_f$) value. After the TLC plates with loaded samples were run through the mobile phase, pNP-cellobiose standards of known concentrations ranging from 0.25 mM to 10 mM were spotted on the TLC plate. The plate was epi-illuminated and directly imaged under UV light at wavelength λ=305 nm to visualize pNP and pNP-containing compounds. The plates were then immediately sprayed with visualization solution containing 0.1% orcinol dye in 10% $H_2SO_4$ in ethyl alcohol, then dried and heated at 100 °C for 15 min to visualize reducing sugars and acid-labile sugars. The plates were then imaged GelDoc EZ imager (BioRad) and spot intensity was quantified using GelDoc EZ imaging software. For quantitative analysis of kinetic assay reaction mixtures, a standard curve was built using the pNPC standards and respective standard spot intensities. The standard curve was then used to estimate the absolute concentration (as pNPC equivalents) of all other hydrolysis or transglycosylation products seen on the TLC plate (e.g., glucose, pNP-glucose, pNP-cellotriose, pNP-cellotetraose, cellobiose, cellotriose, cellotetraose, and cellopentaose). Fractional concentrations for each product reported here was normalized to the total observed residual substrate and formed products concentration for each reaction mixture. Final results reported here for each reaction condition was based on two biological replicates.

***SAXS analysis***: Purified protein samples for SAXS analysis were diluted in 50 mM MES pH 6.5 buffer to get a final concentration of 1 mg/ml and 2.5 mg/ml. The reaction samples were prepared to consist of protein at 1 mg/ml and 2.5 mg/ml and substrate (pNPC) at 10 mM as final concentration in 50 mM MES pH 6.5 buffer. All the samples mixtures were prepared on ice and instantly frozen using liquid nitrogen and shipped on dry ice to Oak Ridge National Laboratory (ORNL) for SAXS measurements. SAXS experiments were carried out on a Rigaku BioSAXS 2000 instrument equipped with a Pilatus 100K detector (Rigaku Americas) at Oak Ridge National Laboratory (ORNL). Silver behenate were used to calibrate for sample-to-detector distance as well as beam center. Sample volumes of ~80-100 µls were loaded into a Julabo temperature-controlled 96-well plate. An automatic sample loader pumped samples from the 96-well plate into the Julabo temperature-controlled flow cell for measurement. The 2D images obtained were reduced using instrument reduction software to 1D curves, $I(Q)$ vs. $Q$, where $Q$ the wave-vector is a function of scattering angle, $2\theta$ and wavelength, $\lambda$. An accompanying buffer was measured for each sample and subsequently, the buffer background was subtracted. The resulting scattering profile representative of the sample was used for data analysis. The pair distance distribution function P(r) of the protein was calculated from the indirect Fourier transform of $I(Q)$ and further a low-resolution volume envelop using a dummy atom model was obtained using the ATSAS package, GNOM and DAMMIN, respectively (2). Ensemble optimization method (EOM) module of ATSAS package was used to generate the model structures. The crystal structures of CelE (PDB ID: 4IM4) and CBM3a (PDB ID: 1NBC) were used as input structures for EOM analysis along with the experimental scattering data.

**Supplementary Text S1.** Protein sequences of all major constructs used in this study.

>> CelE-Wt
GMRDISAIDLVKEIKIGWNLGNTLDAPTETAWGNPRTTKAMIEKVREMGFNAVRVPVTWDTHIGPAPDYKIDEAWLNRVEE
VVNYVLDCGMYAIINVHHDNTWIIPTYANEQRSKEKLVKVWEQIATRFKDYDDHLLFETMNEPREVGSPMEWMGGTYENR
DVINRFNLAVVNTIRASGGNNDKRFILVPTNAATGLDVALNDLVIPNNDSRVIVSIHAYSPYFFAMDVNGTSYWGSDYDKAS
FTSELDAIYNRFVKNGRAVIIGEFGTIDKNNLSSRVAHAEHYAREAVSRGIAVFWWDNGYYNPGDAETYALLNRRNLTWYY
PEIVQALMRGAG

>> CelE-E316A
GMRDISAIDLVKEIKIGWNLGNTLDAPTETAWGNPRTTKAMIEKVREMGFNAVRVPVTWDTHIGPAPDYKIDEAWLNRVEE
VVNYVLDCGMYAIINVHHDNTWIIPTYANEQRSKEKLVKVWEQIATRFKDYDDHLLFETMNEPREVGSPMEWMGGTYENR
DVINRFNLAVVNTIRASGGNNDKRFILVPTNAATGLDVALNDLVIPNNDSRVIVSIHAYSPYFFAMDVNGTSYWGSDYDKAS
FTSELDAIYNRFVKNGRAVIIGAFGTIDKNNLSSRVAHAEHYAREAVSRGIAVFWWDNGYYNPGDAETYALLNRRNLTWYY
PEIVQALMRGAG

>> CelE-E316S
GMRDISAIDLVKEIKIGWNLGNTLDAPTETAWGNPRTTKAMIEKVREMGFNAVRVPVTWDTHIGPAPDYKIDEAWLNRVEE
VVNYVLDCGMYAIINVHHDNTWIIPTYANEQRSKEKLVKVWEQIATRFKDYDDHLLFETMNEPREVGSPMEWMGGTYENR
DVINRFNLAVVNTIRASGGNNDKRFILVPTNAATGLDVALNDLVIPNNDSRVIVSIHAYSPYFFAMDVNGTSYWGSDYDKAS
FTSELDAIYNRFVKNGRAVIIGSFGTIDKNNLSSRVAHAEHYAREAVSRGIAVFWWDNGYYNPGDAETYALLNRRNLTWYY
PEIVQALMRGAG

>> CelE-E316G
GMRDISAIDLVKEIKIGWNLGNTLDAPTETAWGNPRTTKAMIEKVREMGFNAVRVPVTWDTHIGPAPDYKIDEAWLNRVEE
VVNYVLDCGMYAIINVHHDNTWIIPTYANEQRSKEKLVKVWEQIATRFKDYDDHLLFETMNAPREVGSPMEWMGGTYENR
DVINRFNLAVVNTIRASGGNNDKRFILVPTNAATGLDVALNDLVIPNNDSRVIVSIHAYSPYFFAMDVNGTSYWGSDYDKAS
FTSELDAIYNRFVKNGRAVIIGGFGTIDKNNLSSRVAHAEHYAREAVSRGIAVFWWDNGYYNPGDAETYALLNRRNLTWYY
PEIVQALMRGAG

>> CelE-Wt-CBM3a-42aaLinker
GMRDISAIDLVKEIKIGWNLGNTLDAPTETAWGNPRTTKAMIEKVREMGFNAVRVPVTWDTHIGPAPDYKIDEAWLNRVEE
VVNYVLDCGMYAIINVHHDNTWIIPTYANEQRSKEKLVKVWEQIATRFKDYDDHLLFETMNEPREVGSPMEWMGGTYENR
DVINRFNLAVVNTIRASGGNNDKRFILVPTNAATGLDVALNDLVIPNNDSRVIVSIHAYSPYFFAMDVNGTSYWGSDYDKAS
FTSELDAIYNRFVKNGRAVIIGEFGTIDKNNLSSRVAHAEHYAREAVSRGIAVFWWDNGYYNPGDAETYALLNRRNLTWYY
PEIVQALMRGAGVEGLNATPTKGATPTNTATPTKSATATPTRPSVPTNTPTNTPANTPVSGNLKVEFYNSNPSDTTNSINP
QFKVTNTGSSAIDLSKLTLRYYYTVDGQKDQTFWCDHAAIIGSNGSYNGITSNVKGTFVKMSSSTNNADTYLEISFTGGTLE
PGAHVQIQGRFAKNDWSNYTQSNDYSFKSASQFVEWDQVTAYLNGVLVWGKEPG

>> CelE- E316A-CBM3a-42aaLinker
GMRDISAIDLVKEIKIGWNLGNTLDAPTETAWGNPRTTKAMIEKVREMGFNAVRVPVTWDTHIGPAPDYKIDEAWLNRVEE
VVNYVLDCGMYAIINVHHDNTWIIPTYANEQRSKEKLVKVWEQIATRFKDYDDHLLFETMNEPREVGSPMEWMGGTYENR
DVINRFNLAVVNTIRASGGNNDKRFILVPTNAATGLDVALNDLVIPNNDSRVIVSIHAYSPYFFAMDVNGTSYWGSDYDKAS
FTSELDAIYNRFVKNGRAVIIGAFGTIDKNNLSSRVAHAEHYAREAVSRGIAVFWWDNGYYNPGDAETYALLNRRNLTWYY
PEIVQALMRGAGVEGLNATPTKGATPTNTATPTKSATATPTRPSVPTNTPTNTPANTPVSGNLKVEFYNSNPSDTTNSINP
QFKVTNTGSSAIDLSKLTLRYYYTVDGQKDQTFWCDHAAIIGSNGSYNGITSNVKGTFVKMSSSTNNADTYLEISFTGGTLE
PGAHVQIQGRFAKNDWSNYTQSNDYSFKSASQFVEWDQVTAYLNGVLVWGKEPG

>> CelE- E316S-CBM3a-42aaLinker
GMRDISAIDLVKEIKIGWNLGNTLDAPTETAWGNPRTTKAMIEKVREMGFNAVRVPVTWDTHIGPAPDYKIDEAWLNRVEE
VVNYVLDCGMYAIINVHHDNTWIIPTYANEQRSKEKLVKVWEQIATRFKDYDDHLLFETMNEPREVGSPMEWMGGTYENR
DVINRFNLAVVNTIRASGGNNDKRFILVPTNAATGLDVALNDLVIPNNDSRVIVSIHAYSPYFFAMDVNGTSYWGSDYDKAS
FTSELDAIYNRFVKNGRAVIIGSFGTIDKNNLSSRVAHAEHYAREAVSRGIAVFWWDNGYYNPGDAETYALLNRRNLTWYY
PEIVQALMRGAGVEGLNATPTKGATPTNTATPTKSATATPTRPSVPTNTPTNTPANTPVSGNLKVEFYNSNPSDTTNSINP
QFKVTNTGSSAIDLSKLTLRYYYTVDGQKDQTFWCDHAAIIGSNGSYNGITSNVKGTFVKMSSSTNNADTYLEISFTGGTLE
PGAHVQIQGRFAKNDWSNYTQSNDYSFKSASQFVEWDQVTAYLNGVLVWGKEPG

>> CelE- E316G-CBM3a-42aaLinker

GMRDISAIDLVKEIKIGWNLGNTLDAPTETAWGNPRTTKAMIEKVREMGFNAVRVPVTWDTHIGPAPDYKIDEAWLNRVEE
VVNYVLDCGMYAIINVHHDNTWIIPTYANEQRSKEKLVKVWEQIATRFKDYDDHLLFETMNEPREVGSPMEWMGGTYENR
DVINRFNLAVVNTIRASGGNNDKRFILVPTNAATGLDVALNDLVIPNNDSRVIVSIHAYSPYFFAMDVNGTSYWGSDYDKAS
FTSELDAIYNRFVKNGRAVIIGGFGTIDKNNLSSRVAHAEHYAREAVSRGIAVFWWDNGYYNPGDAETYALLNRRNLTWYY
PEIVQALMRGAGVEGLNATPTKGATPTNTATPTKSATATPTRPSVPTNTPTNTPANTPVSGNLKVEFYNSNPSDTTNSINP
QFKVTNTGSSAIDLSKLTLRYYYTVDGQKDQTFWCDHAAIIGSNGSYNGITSNVKGTFVKMSSSTNNADTYLEISFTGGTLE
PGAHVQIQGRFAKNDWSNYTQSNDYSFKSASQFVEWDQVTAYLNGVLVWGKEPG

>> CelE-Wt-CBM1-42aaLinker
GMRDISAIDLVKEIKIGWNLGNTLDAPTETAWGNPRTTKAMIEKVREMGFNAVRVPVTWDTHIGPAPDYKIDEAWLNRVEE
VVNYVLDCGMYAIINVHHDNTWIIPTYANEQRSKEKLVKVWEQIATRFKDYDDHLLFETMNEPREVGSPMEWMGGTYENR
DVINRFNLAVVNTIRASGGNNDKRFILVPTNAATGLDVALNDLVIPNNDSRVIVSIHAYSPYFFAMDVNGTSYWGSDYDKAS
FTSELDAIYNRFVKNGRAVIIGEFGTIDKNNLSSRVAHAEHYAREAVSRGIAVFWWDNGYYNPGDAETYALLNRRNLTWYY
PEIVQALMRGAGVEGLNATPTKGATPTNTATPTKSATATPTRPSVPTNTPTNTPANTLKPGPTQSHYGQCGGIGYSGPTV
CASGTTCQVLNPYYSQCL

>> CelE-Wt-CBM17-42aaLinker
GMRDISAIDLVKEIKIGWNLGNTLDAPTETAWGNPRTTKAMIEKVREMGFNAVRVPVTWDTHIGPAPDYKIDEAWLNRVEE
VVNYVLDCGMYAIINVHHDNTWIIPTYANEQRSKEKLVKVWEQIATRFKDYDDHLLFETMNEPREVGSPMEWMGGTYENR
DVINRFNLAVVNTIRASGGNNDKRFILVPTNAATGLDVALNDLVIPNNDSRVIVSIHAYSPYFFAMDVNGTSYWGSDYDKAS
FTSELDAIYNRFVKNGRAVIIGEFGTIDKNNLSSRVAHAEHYAREAVSRGIAVFWWDNGYYNPGDAETYALLNRRNLTWYY
PEIVQALMRGAGVEGLNATPTKGATPTNTATPTKSATATPTRPSVPTNTPTNTPANTLKSQPTAPKDFSSGFWDFNDGTT
QGFGVNPDSPITAINVENANNALKISNLNSKGSNDLSEGNFWANVRISADIWGQSINIYGDTKLTMDVIAPTPVNVSIAAIPQ
SSTHGWGNPTRAIRVWTNNFVAQTDGTYKATLTISTNDSPNFNTIATDAADSVVTNMILFVGSNSDNISLDNIKFTK

>> CelE-Wt-CBM3a- 6aaLinker
GMRDISAIDLVKEIKIGWNLGNTLDAPTETAWGNPRTTKAMIEKVREMGFNAVRVPVTWDTHIGPAPDYKIDEAWLNRVEE
VVNYVLDCGMYAIINVHHDNTWIIPTYANEQRSKEKLVKVWEQIATRFKDYDDHLLFETMNEPREVGSPMEWMGGTYENR
DVINRFNLAVVNTIRASGGNNDKRFILVPTNAATGLDVALNDLVIPNNDSRVIVSIHAYSPYFFAMDVNGTSYWGSDYDKAS
FTSELDAIYNRFVKNGRAVIIGEFGTIDKNNLSSRVAHAEHYAREAVSRGIAVFWWDNGYYNPGDAETYALLNRRNLTWYY
PEIVQALMRGAGVEGLNATPTKVSGNLKVEFYNSNPSDTTNSINPQFKVTNTGSSAIDLSKLTLRYYYTVDGQKDQTFWCD
HAAIIGSNGSYNGITSNVKGTFVKMSSSTNNADTYLEISFTGGTLEPGAHVQIQGRFAKNDWSNYTQSNDYSFKSASQFVE
WDQVTAYLNGVLVWGKEPG

>> CelE-Wt-CBM3a-11aaLinker
GMRDISAIDLVKEIKIGWNLGNTLDAPTETAWGNPRTTKAMIEKVREMGFNAVRVPVTWDTHIGPAPDYKIDEAWLNRVEE
VVNYVLDCGMYAIINVHHDNTWIIPTYANEQRSKEKLVKVWEQIATRFKDYDDHLLFETMNEPREVGSPMEWMGGTYENR
DVINRFNLAVVNTIRASGGNNDKRFILVPTNAATGLDVALNDLVIPNNDSRVIVSIHAYSPYFFAMDVNGTSYWGSDYDKAS
FTSELDAIYNRFVKNGRAVIIGEFGTIDKNNLSSRVAHAEHYAREAVSRGIAVFWWDNGYYNPGDAETYALLNRRNLTWYY
PEIVQALMRGAGVEGLNATPTKGATPTVSGNLKVEFYNSNPSDTTNSINPQFKVTNTGSSAIDLSKLTLRYYYTVDGQKDQ
TFWCDHAAIIGSNGSYNGITSNVKGTFVKMSSSTNNADTYLEISFTGGTLEPGAHVQIQGRFAKNDWSNYTQSNDYSFKSA
SQFVEWDQVTAYLNGVLVWGKEPG

>> CelE-Wt-CBM3a-21aaLinker
GMRDISAIDLVKEIKIGWNLGNTLDAPTETAWGNPRTTKAMIEKVREMGFNAVRVPVTWDTHIGPAPDYKIDEAWLNRVEE
VVNYVLDCGMYAIINVHHDNTWIIPTYANEQRSKEKLVKVWEQIATRFKDYDDHLLFETMNEPREVGSPMEWMGGTYENR
DVINRFNLAVVNTIRASGGNNDKRFILVPTNAATGLDVALNDLVIPNNDSRVIVSIHAYSPYFFAMDVNGTSYWGSDYDKAS
FTSELDAIYNRFVKNGRAVIIGEFGTIDKNNLSSRVAHAEHYAREAVSRGIAVFWWDNGYYNPGDAETYALLNRRNLTWYY
PEIVQALMRGAGVEGLNATPTKGATPTNTATPTKSATVSGNLKVEFYNSNPSDTTNSINPQFKVTNTGSSAIDLSKLTLRYY
YTVDGQKDQTFWCDHAAIIGSNGSYNGITSNVKGTFVKMSSSTNNADTYLEISFTGGTLEPGAHVQIQGRFAKNDWSNYT
QSNDYSFKSASQFVEWDQVTAYLNGVLVWGKEPG

>> CelE-Wt-CBM3a-Flex1
GMRDISAIDLVKEIKIGWNLGNTLDAPTETAWGNPRTTKAMIEKVREMGFNAVRVPVTWDTHIGPAPDYKIDEAWLNRVEE
VVNYVLDCGMYAIINVHHDNTWIIPTYANEQRSKEKLVKVWEQIATRFKDYDDHLLFETMNEPREVGSPMEWMGGTYENR
DVINRFNLAVVNTIRASGGNNDKRFILVPTNAATGLDVALNDLVIPNNDSRVIVSIHAYSPYFFAMDVNGTSYWGSDYDKAS
FTSELDAIYNRFVKNGRAVIIGEFGTIDKNNLSSRVAHAEHYAREAVSRGIAVFWWDNGYYNPGDAETYALLNRRNLTWYY
PEIVQALMRGAGVEGLNATGTKGATGTNTATGTKSATATGTRGSVGTNTGTNTGANTGVSGNLKVEFYNSNPSDTTNSIN

PQFKVTNTGSSAIDLSKLTLRYYYTVDGQKDQTFWCDHAAIIGSNGSYNGITSNVKGTFVKMSSSTNNADTYLEISFTGGTL
EPGAHVQIQGRFAKNDWSNYTQSNDYSFKSASQFVEWDQVTAYLNGVLVWGKEPG

>> CelE-Wt-CBM3a-Flex2
GMRDISAIDLVKEIKIGWNLGNTLDAPTETAWGNPRTTKAMIEKVREMGFNAVRVPVTWDTHIGPAPDYKIDEAWLNRVEE
VVNYVLDCGMYAIINVHHDNTWIIPTYANEQRSKEKLVKVWEQIATRFKDYDDHLLFETMNEPREVGSPMEWMGGTYENR
DVINRFNLAVVNTIRASGGNNDKRFILVPTNAATGLDVALNDLVIPNNDSRVIVSIHAYSPYFFAMDVNGTSYWGSDYDKAS
FTSELDAIYNRFVKNGRAVIIGEFGTIDKNNLSSRVAHAEHYAREAVSRGIAVFWWDNGYYNPGDAETYALLNRRNLTWYY
PEIVQALMRGAGVEGLNASGGKGATGGNTAGGTKSATAGGSRGSVGGNSGTNGGANGGVSGNLKVEFYNSNPSDTTN
SINPQFKVTNTGSSAIDLSKLTLRYYYTVDGQKDQTFWCDHAAIIGSNGSYNGITSNVKGTFVKMSSSTNNADTYLEISFTG
GTLEPGAHVQIQGRFAKNDWSNYTQSNDYSFKSASQFVEWDQVTAYLNGVLVWGKEPG

>> CelE-Wt-CBM3a-Rig1
GMRDISAIDLVKEIKIGWNLGNTLDAPTETAWGNPRTTKAMIEKVREMGFNAVRVPVTWDTHIGPAPDYKIDEAWLNRVEE
VVNYVLDCGMYAIINVHHDNTWIIPTYANEQRSKEKLVKVWEQIATRFKDYDDHLLFETMNEPREVGSPMEWMGGTYENR
DVINRFNLAVVNTIRASGGNNDKRFILVPTNAATGLDVALNDLVIPNNDSRVIVSIHAYSPYFFAMDVNGTSYWGSDYDKAS
FTSELDAIYNRFVKNGRAVIIGEFGTIDKNNLSSRVAHAEHYAREAVSRGIAVFWWDNGYYNPGDAETYALLNRRNLTWYY
PEIVQALMRGAGVEGLNATPTKPATPTNTPTPTKPATATPTRPPVPTNTPTNTPANTPVSGNLKVEFYNSNPSDTTNSINP
QFKVTNTGSSAIDLSKLTLRYYYTVDGQKDQTFWCDHAAIIGSNGSYNGITSNVKGTFVKMSSSTNNADTYLEISFTGGTLE
PGAHVQIQGRFAKNDWSNYTQSNDYSFKSASQFVEWDQVTAYLNGVLVWGKEPG

>> CelE-Wt-CBM3a-Rig2
GMRDISAIDLVKEIKIGWNLGNTLDAPTETAWGNPRTTKAMIEKVREMGFNAVRVPVTWDTHIGPAPDYKIDEAWLNRVEE
VVNYVLDCGMYAIINVHHDNTWIIPTYANEQRSKEKLVKVWEQIATRFKDYDDHLLFETMNEPREVGSPMEWMGGTYENR
DVINRFNLAVVNTIRASGGNNDKRFILVPTNAATGLDVALNDLVIPNNDSRVIVSIHAYSPYFFAMDVNGTSYWGSDYDKAS
FTSELDAIYNRFVKNGRAVIIGEFGTIDKNNLSSRVAHAEHYAREAVSRGIAVFWWDNGYYNPGDAETYALLNRRNLTWYY
PEIVQALMRGAGVEGLGGSAEAAAKAAEAAAKAAEAAAKAAEAAAKAAEAAAKASGGVSGNLKVEFYNSNPSDTTNSINP
QFKVTNTGSSAIDLSKLTLRYYYTVDGQKDQTFWCDHAAIIGSNGSYNGITSNVKGTFVKMSSSTNNADTYLEISFTGGTLE
PGAHVQIQGRFAKNDWSNYTQSNDYSFKSASQFVEWDQVTAYLNGVLVWGKEPG

>> CelE-Wt-CBM3a-Y458A
GMRDISAIDLVKEIKIGWNLGNTLDAPTETAWGNPRTTKAMIEKVREMGFNAVRVPVTWDTHIGPAPDYKIDEAWLNRVEE
VVNYVLDCGMYAIINVHHDNTWIIPTYANEQRSKEKLVKVWEQIATRFKDYDDHLLFETMNEPREVGSPMEWMGGTYENR
DVINRFNLAVVNTIRASGGNNDKRFILVPTNAATGLDVALNDLVIPNNDSRVIVSIHAYSPYFFAMDVNGTSYWGSDYDKAS
FTSELDAIYNRFVKNGRAVIIGEFGTIDKNNLSSRVAHAEHYAREAVSRGIAVFWWDNGYYNPGDAETYALLNRRNLTWYY
PEIVQALMRGAGVEGLNATPTKGATPTNTATPTKSATATPTRPSVPTNTPTNTPANTPVSGNLKVEFYNSNPSDTTNSINP
QFKVTNTGSSAIDLSKLTLRYYYTVDGQKDQTFWCDHAAIIGSNGSYNGITSNVKGTFVKMSSSTNNADTALEISFTGGTLE
PGAHVQIQGRFAKNDWSNYTQSNDYSFKSASQFVEWDQVTAYLNGVLVWGKEPG

>> CelE-Wt-CBM3a-E521A
GMRDISAIDLVKEIKIGWNLGNTLDAPTETAWGNPRTTKAMIEKVREMGFNAVRVPVTWDTHIGPAPDYKIDEAWLNRVEE
VVNYVLDCGMYAIINVHHDNTWIIPTYANEQRSKEKLVKVWEQIATRFKDYDDHLLFETMNEPREVGSPMEWMGGTYENR
DVINRFNLAVVNTIRASGGNNDKRFILVPTNAATGLDVALNDLVIPNNDSRVIVSIHAYSPYFFAMDVNGTSYWGSDYDKAS
FTSELDAIYNRFVKNGRAVIIGEFGTIDKNNLSSRVAHAEHYAREAVSRGIAVFWWDNGYYNPGDAETYALLNRRNLTWYY
PEIVQALMRGAGVEGLNATPTKGATPTNTATPTKSATATPTRPSVPTNTPTNTPANTPVSGNLKVEFYNSNPSDTTNSINP
QFKVTNTGSSAIDLSKLTLRYYYTVDGQKDQTFWCDHAAIIGSNGSYNGITSNVKGTFVKMSSSTNNADTYLEISFTGGTLE
PGAHVQIQGRFAKNDWSNYTQSNDYSFKSASQFVEWDQVTAYLNGVLVWGKAPG

>> CelE-Wt-CBM3a-R407A
GMRDISAIDLVKEIKIGWNLGNTLDAPTETAWGNPRTTKAMIEKVREMGFNAVRVPVTWDTHIGPAPDYKIDEAWLNRVEE
VVNYVLDCGMYAIINVHHDNTWIIPTYANEQRSKEKLVKVWEQIATRFKDYDDHLLFETMNEPREVGSPMEWMGGTYENR
DVINRFNLAVVNTIRASGGNNDKRFILVPTNAATGLDVALNDLVIPNNDSRVIVSIHAYSPYFFAMDVNGTSYWGSDYDKAS
FTSELDAIYNRFVKNGRAVIIGEFGTIDKNNLSSRVAHAEHYAREAVSRGIAVFWWDNGYYNPGDAETYALLNRRNLTWYY
PEIVQALMRGAGVEGLNATPTKGATPTNTATPTKSATATPTRPSVPTNTPTNTPANTPVSGNLKVEFYNSNPSDTTNSINP
QFKVTNTGSSAIDLSKLTLAYYYTVDGQKDQTFWCDHAAIIGSNGSYNGITSNVKGTFVKMSSSTNNADTYLEISFTGGTLE
PGAHVQIQGRFAKNDWSNYTQSNDYSFKSASQFVEWDQVTAYLNGVLVWGKEPG

>> CelE-Wt-CBM3a-T509A

GMRDISAIDLVKEIKIGWNLGNTLDAPTETAWGNPRTTKAMIEKVREMGFNAVRVPVTWDTHIGPAPDYKIDEAWLNRVEE
VVNYVLDCGMYAIINVHHDNTWIIPTYANEQRSKEKLVKVWEQIATRFKDYDDHLLFETMNEPREVGSPMEWMGGTYENR
DVINRFNLAVVNTIRASGGNNDKRFILVPTNAATGLDVALNDLVIPNNDSRVIVSIHAYSPYFFAMDVNGTSYWGSDYDKAS
FTSELDAIYNRFVKNGRAVIIGEFGTIDKNNLSSRVAHAEHYAREAVSRGIAVFWWDNGYYNPGDAETYALLNRRNLTWYY
PEIVQALMRGAGVEGLNATPTKGATPTNTATPTKSATATPTRPSVPTNTPTNTPANTPVSGNLKVEFYNSNPSDTTNSINP
QFKVTNTGSSAIDLSKLTLRYYYTVDGQKDQTFWCDHAAIIGSNGSYNGITSNVKGTFVKMSSSTNNADTYLEISFTGGTLE
PGAHVQIQGRFAKNDWSNYTQSNDYSFKSASQFVEWDQVAAYLNGVLVWGKEPG

>> CelE-Wt-CBM3a-R409A
GMRDISAIDLVKEIKIGWNLGNTLDAPTETAWGNPRTTKAMIEKVREMGFNAVRVPVTWDTHIGPAPDYKIDEAWLNRVEE
VVNYVLDCGMYAIINVHHDNTWIIPTYANEQRSKEKLVKVWEQIATRFKDYDDHLLFETMNEPREVGSPMEWMGGTYENR
DVINRFNLAVVNTIRASGGNNDKRFILVPTNAATGLDVALNDLVIPNNDSRVIVSIHAYSPYFFAMDVNGTSYWGSDYDKAS
FTSELDAIYNRFVKNGRAVIIGEFGTIDKNNLSSRVAHAEHYAREAVSRGIAVFWWDNGYYNPGDAETYALLNRRNLTWYY
PEIVQALMRGAGVEGLNATPTKGATPTNTATPTKSATATPTRPSVPTNTPTNTPANTPVSGNLKVEFYNSNPSDTTNSINP
QFKVTNTGSSAIDLSKLTLRYAYTVDGQKDQTFWCDHAAIIGSNGSYNGITSNVKGTFVKMSSSTNNADTYLEISFTGGTLE
PGAHVQIQGRFAKNDWSNYTQSNDYSFKSASQFVEWDQVTAYLNGVLVWGKEPG

>> CelE-Y273A-CBM3a
GMRDISAIDLVKEIKIGWNLGNTLDAPTETAWGNPRTTKAMIEKVREMGFNAVRVPVTWDTHIGPAPDYKIDEAWLNRVEE
VVNYVLDCGMYAIINVHHDNTWIIPTYANEQRSKEKLVKVWEQIATRFKDYDDHLLFETMNEPREVGSPMEWMGGTYENR
DVINRFNLAVVNTIRASGGNNDKRFILVPTNAATGLDVALNDLVIPNNDSRVIVSIHAYSPAFFAMDVNGTSYWGSDYDKAS
FTSELDAIYNRFVKNGRAVIIGEFGTIDKNNLSSRVAHAEHYAREAVSRGIAVFWWDNGYYNPGDAETYALLNRRNLTWYY
PEIVQALMRGAGVEGLNATPTKGATPTNTATPTKSATATPTRPSVPTNTPTNTPANTPVSGNLKVEFYNSNPSDTTNSINP
QFKVTNTGSSAIDLSKLTLRYYYTVDGQKDQTFWCDHAAIIGSNGSYNGITSNVKGTFVKMSSSTNNADTYLEISFTGGTLE
PGAHVQIQGRFAKNDWSNYTQSNDYSFKSASQFVEWDQVTAYLNGVLVWGKEPG

>> CelE-Y270A-CBM3a
GMRDISAIDLVKEIKIGWNLGNTLDAPTETAWGNPRTTKAMIEKVREMGFNAVRVPVTWDTHIGPAPDYKIDEAWLNRVEE
VVNYVLDCGMYAIINVHHDNTWIIPTYANEQRSKEKLVKVWEQIATRFKDYDDHLLFETMNEPREVGSPMEWMGGTYENR
DVINRFNLAVVNTIRASGGNNDKRFILVPTNAATGLDVALNDLVIPNNDSRVIVSIHAASPYFFAMDVNGTSYWGSDYDKAS
FTSELDAIYNRFVKNGRAVIIGEFGTIDKNNLSSRVAHAEHYAREAVSRGIAVFWWDNGYYNPGDAETYALLNRRNLTWYY
PEIVQALMRGAGVEGLNATPTKGATPTNTATPTKSATATPTRPSVPTNTPTNTPANTPVSGNLKVEFYNSNPSDTTNSINP
QFKVTNTGSSAIDLSKLTLRYYYTVDGQKDQTFWCDHAAIIGSNGSYNGITSNVKGTFVKMSSSTNNADTYLEISFTGGTLE
PGAHVQIQGRFAKNDWSNYTQSNDYSFKSASQFVEWDQVTAYLNGVLVWGKEPG

>> CelE-H268A-CBM3a
GMRDISAIDLVKEIKIGWNLGNTLDAPTETAWGNPRTTKAMIEKVREMGFNAVRVPVTWDTHIGPAPDYKIDEAWLNRVEE
VVNYVLDCGMYAIINVHHDNTWIIPTYANEQRSKEKLVKVWEQIATRFKDYDDHLLFETMNEPREVGSPMEWMGGTYENR
DVINRFNLAVVNTIRASGGNNDKRFILVPTNAATGLDVALNDLVIPNNDSRVIVSIAAYSPYFFAMDVNGTSYWGSDYDKAS
FTSELDAIYNRFVKNGRAVIIGEFGTIDKNNLSSRVAHAEHYAREAVSRGIAVFWWDNGYYNPGDAETYALLNRRNLTWYY
PEIVQALMRGAGVEGLNATPTKGATPTNTATPTKSATATPTRPSVPTNTPTNTPANTPVSGNLKVEFYNSNPSDTTNSINP
QFKVTNTGSSAIDLSKLTLRYYYTVDGQKDQTFWCDHAAIIGSNGSYNGITSNVKGTFVKMSSSTNNADTYLEISFTGGTLE
PGAHVQIQGRFAKNDWSNYTQSNDYSFKSASQFVEWDQVTAYLNGVLVWGKEPG

>> CelE-W203A-CBM3a
GMRDISAIDLVKEIKIGWNLGNTLDAPTETAWGNPRTTKAMIEKVREMGFNAVRVPVTWDTHIGPAPDYKIDEAWLNRVEE
VVNYVLDCGMYAIINVHHDNTWIIPTYANEQRSKEKLVKVWEQIATRFKDYDDHLLFETMNEPREVGSPMEAMGGTYENR
DVINRFNLAVVNTIRASGGNNDKRFILVPTNAATGLDVALNDLVIPNNDSRVIVSIHAYSPYFFAMDVNGTSYWGSDYDKAS
FTSELDAIYNRFVKNGRAVIIGEFGTIDKNNLSSRVAHAEHYAREAVSRGIAVFWWDNGYYNPGDAETYALLNRRNLTWYY
PEIVQALMRGAGVEGLNATPTKGATPTNTATPTKSATATPTRPSVPTNTPTNTPANTPVSGNLKVEFYNSNPSDTTNSINP
QFKVTNTGSSAIDLSKLTLRYYYTVDGQKDQTFWCDHAAIIGSNGSYNGITSNVKGTFVKMSSSTNNADTYLEISFTGGTLE
PGAHVQIQGRFAKNDWSNYTQSNDYSFKSASQFVEWDQVTAYLNGVLVWGKEPG

>> CelE-W349A-CBM3a
GMRDISAIDLVKEIKIGWNLGNTLDAPTETAWGNPRTTKAMIEKVREMGFNAVRVPVTWDTHIGPAPDYKIDEAWLNRVEE
VVNYVLDCGMYAIINVHHDNTWIIPTYANEQRSKEKLVKVWEQIATRFKDYDDHLLFETMNEPREVGSPMEWMGGTYENR
DVINRFNLAVVNTIRASGGNNDKRFILVPTNAATGLDVALNDLVIPNNDSRVIVSIHAYSPYFFAMDVNGTSYWGSDYDKAS
FTSELDAIYNRFVKNGRAVIIGEFGTIDKNNLSSRVAHAEHYAREAVSRGIAVFWADNGYYNPGDAETYALLNRRNLTWYY

PEIVQALMRGAGVEGLNATPTKGATPTNTATPTKSATATPTRPSVPTNTPTNTPANTPVSGNLKVEFYNSNPSDTTNSINP
QFKVTNTGSSAIDLSKLTLRYYYTVDGQKDQTFWCDHAAIIGSNGSYNGITSNVKGTFVKMSSSTNNADTYLEISFTGGTLE
PGAHVQIQGRFAKNDWSNYTQSNDYSFKSASQFVEWDQVTAYLNGVLVWGKEPG

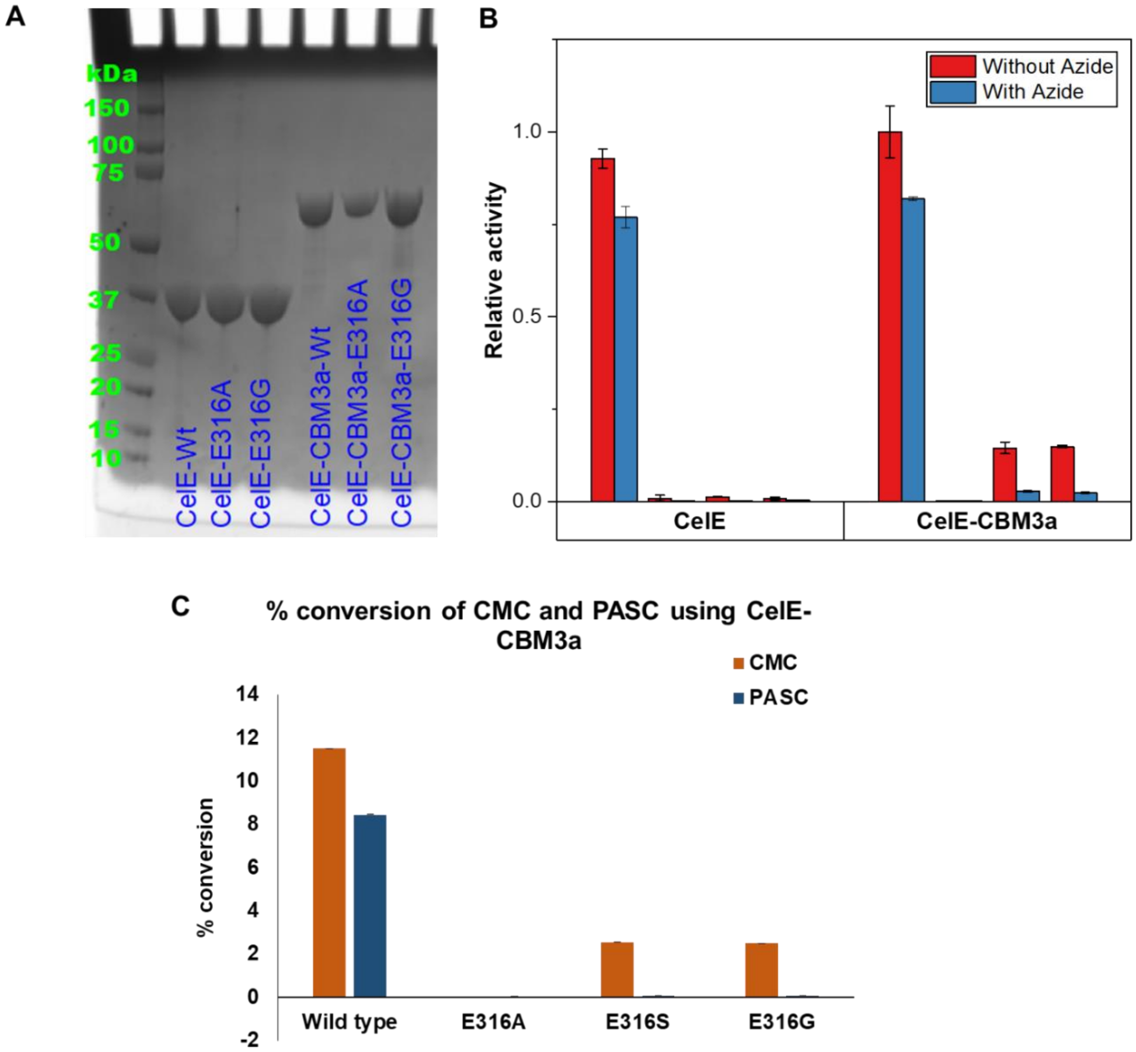


**Fig. S1. SDS-PAGE for purified CelE and CelE-CBM3a wild-type and several mutant enzyme constructs (A), chemical rescue assay (w/wo sodium azide as exogenously added nucleophile) using pNP-cellobiose as substrate with purified enzymes measuring pNP release activity (B), and hydrolytic activity on carboxymethyl cellulose (CMC) and phosphoric acid swollen cellulose (PASC) with purified enzymes (C).** In (B), the 4 sets of bar graphs correspond to wild-type, E316A, E316S, and E316G mutants either without (left) or without (right) a tethered CBM3a domain, respectively. In (C), 10 µg of each protein was incubated with 1 mg of CMC or PASC and the reaction was run for 4 hours at 60°C. DNS assay was performed on the supernatant of the reaction mixtures to measure the amount of hydrolyzed soluble sugars and their respective percent conversion was calculated.

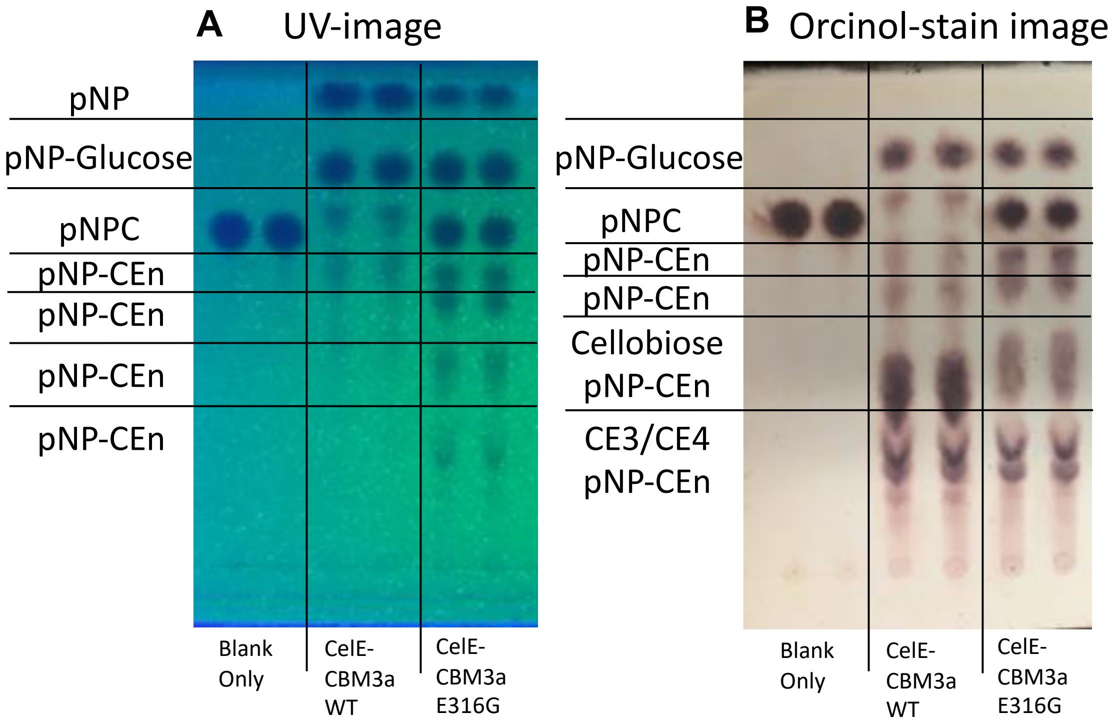


**Fig. S2. Catalytic activity of wild-type CelE-CBM3a and its E316G mutant at high pNP-Cellobiose (pNPC) substrate loadings to clearly visualize pNP-based oligosaccharides formed during the reaction.** Here, 2 nanomoles of each purified protein was incubated along with 12 µmoles of pNPC and the reaction was run for 4 h, respectively, at 60°C reaction temperature. All reactions were run in duplicate as shown below. UV image (A) and orcinol-stained (B) images of the reaction mixtures run on the TLC plates are shown here. Here, CE-‘n’ stands for cello-oligosaccharides with degree of polymerization ‘n’ greater than 2.

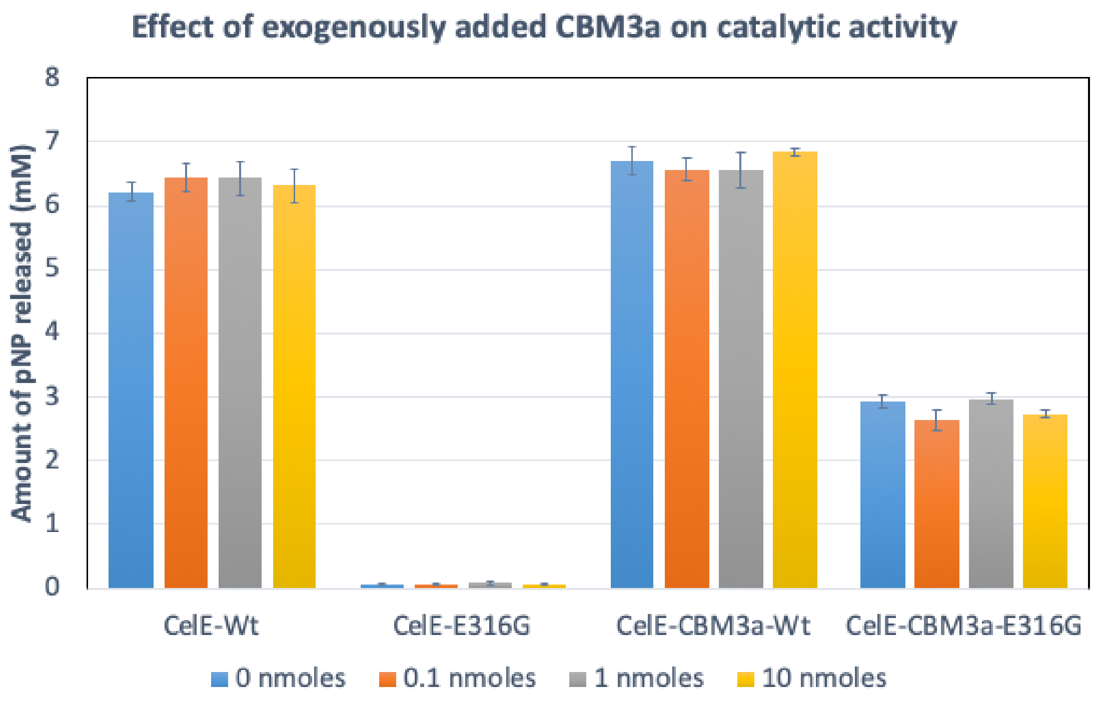


**Fig. S3. Intermolecular interactions of exogenously added CBM3a domain along with CelE-E316G catalytic nucleophile mutant does not increase transglycosylation activity.** Here, 1 nanomoles of each purified protein was incubated along with 1.5 µmoles of pNP-Cellobiose (pNPC) and the reaction was run for 4 h at 60°C reaction temperature. Different concentrations of purified CBM3a domain alone was spiked into the reaction mixture at 0.1-10 times the relative molar ratio of the added CelE or CelE-CBM enzyme construct. All reactions were run in duplicate with error bars representing 1σ from reported mean value. Total pNP released after 4 h reaction time is shown here.

**A**

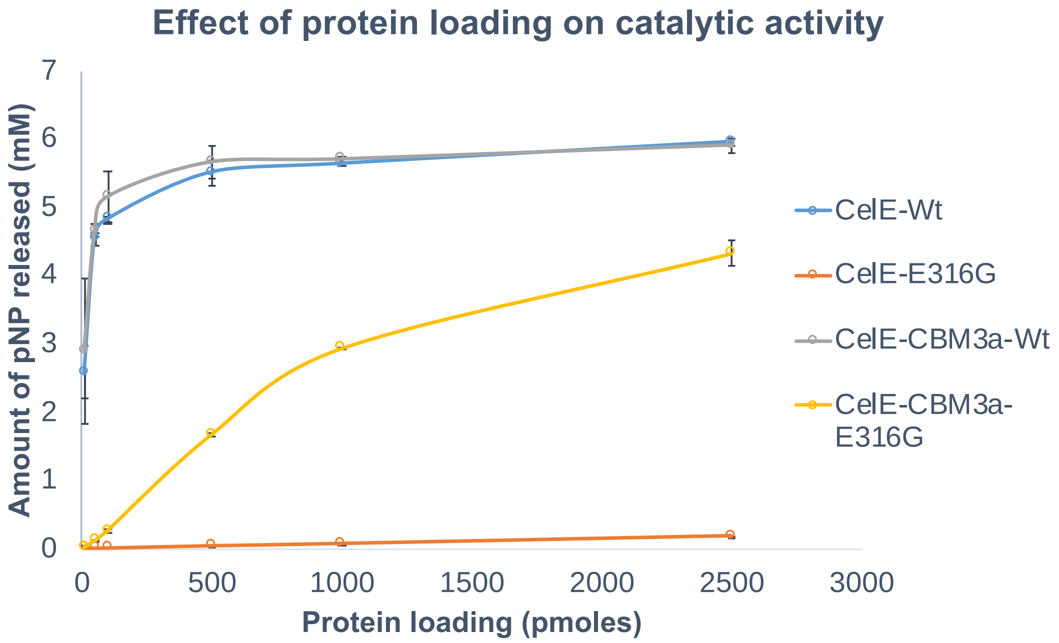


**B**

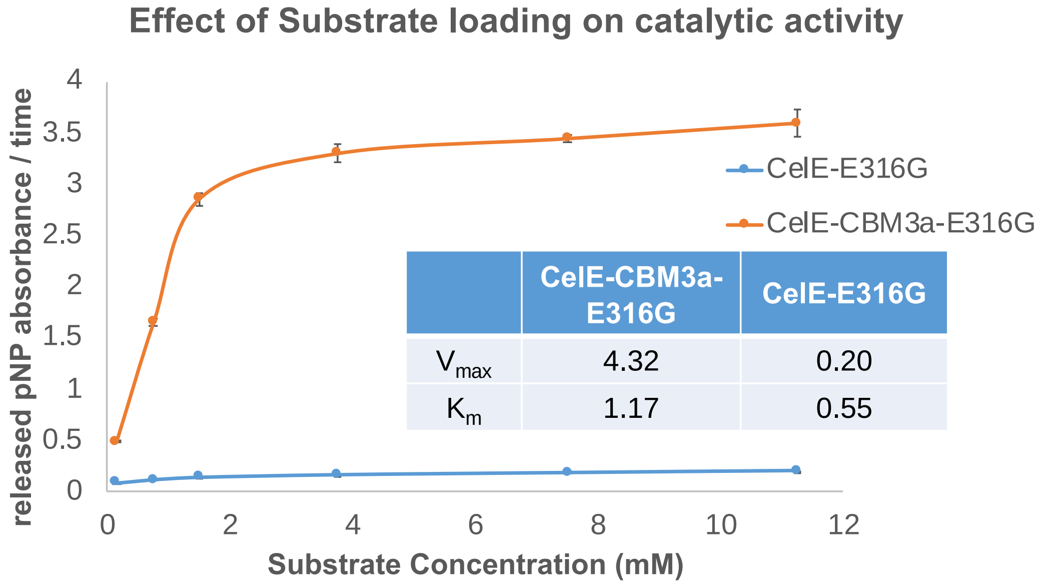


| | CelE-CBM3a-E316G | CelE-E316G |
|---|---|---|
| $V_{max}$ | 4.32 | 0.20 |
| $K_m$ | 1.17 | 0.55 |

**Fig. S4. Effect of protein concentration and substrate concentration on transglycosylation catalytic activity of CelE and CelE-CBM3a based enzyme constructs.** A) 10-to-2500 picomoles of each purified protein was incubated along with fixed 1.5 µmoles of pNP-Cellobiose (pNPC) and the reaction was run for 4 h at 60°C reaction temperature. B) 1000 picomoles of each purified protein was incubated along with 0.15-to-11.25 µmoles of pNP-Cellobiose (pNPC) and the reaction was run for 3 h at 60°C reaction temperature. Here, $K_m$ and $V_{max}$ was calculated by fitting of standard Michaelis-Menten model to data shown in (B). All reactions were run in triplicates (to measure pNP release activity) with error bars representing 1σ from reported mean value.

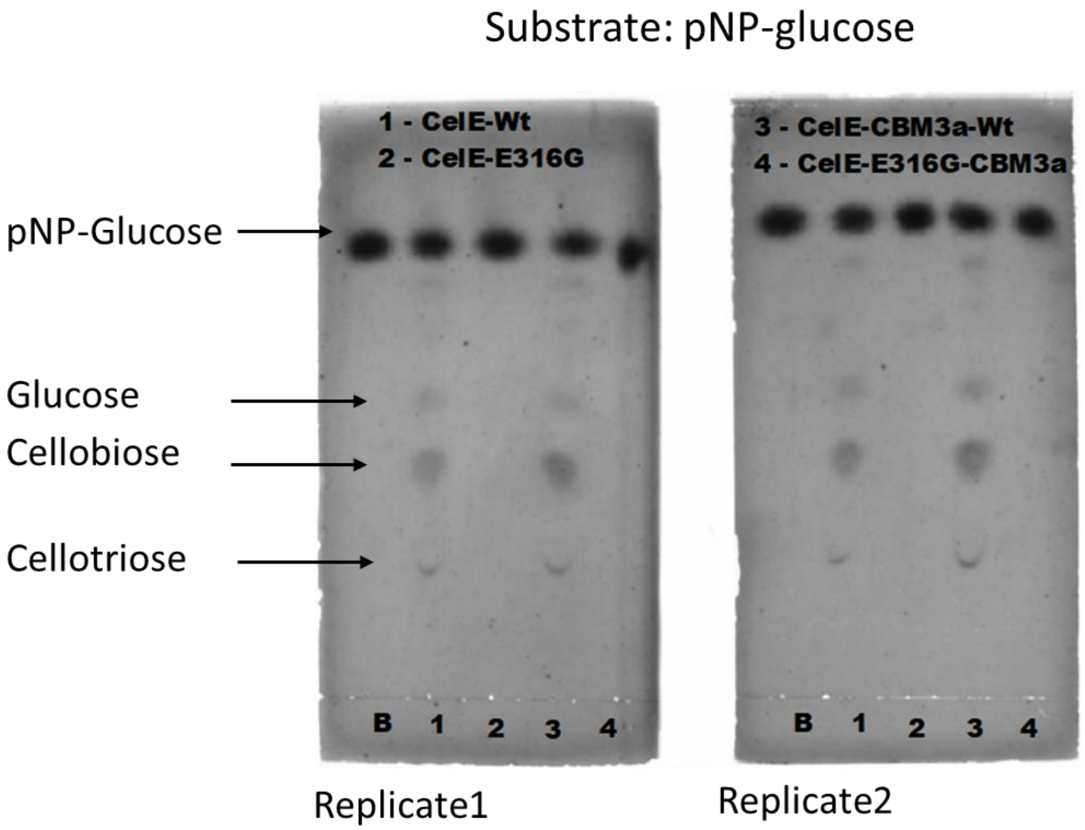


**Fig. S5. Effect of CelE and CelE-CBM3a based constructs catalytic activity on pNP-glucose as starting substrate.** Here, 1 nanomoles of protein was incubated along with fixed 0.5 µmoles of pNP-glucose (pNPG) for 18 h at 60°C reaction temperature. All reactions were run in duplicate. Orcinol-stained images of the reaction mixtures run on the TLC plates are shown here. Here, B- stands for substrate blank (pNPG) alone. Lane 1 – CelE-Wt, Lane – CelE-E316G, Lane 3- CelE-CBM3a-Wt, and Lane 4 – CelE-E316G-CBM3a.

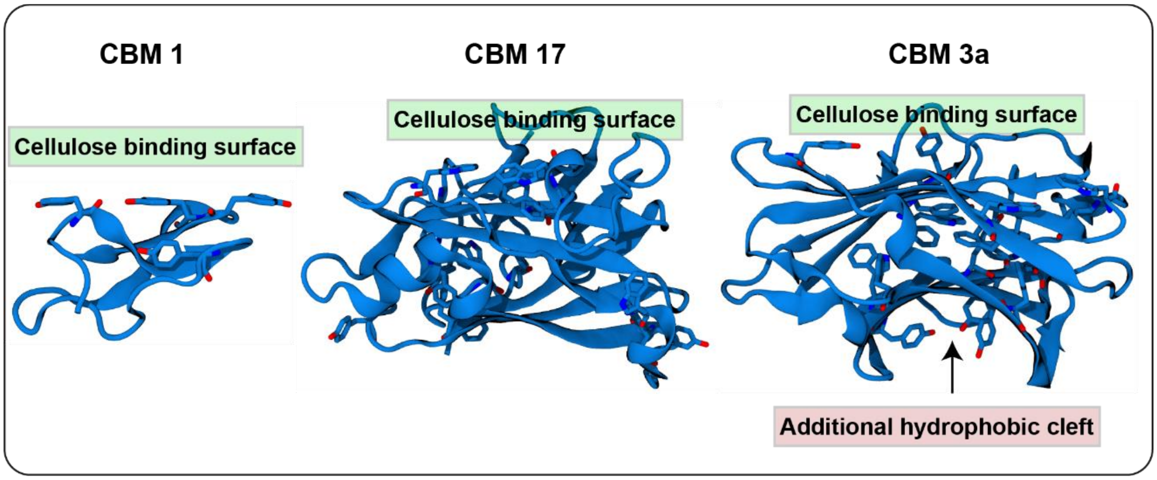


**Fig. S6. Comparing crystal structures of CBM1, CBM17, and CBM3a.** The classical cellulose binding sites/surfaces that could target binding to cellobiose (or pNP-cellobiose) are shown here. Additionally, the hydrophobic cleft in CBM3a on the opposite side of the classical binding surface is also illustrated here. The additional hydrophobic cleft is hypothesized to interact with the pNP-cellobiose substrate to facilitate transglycosylation in CelE-E316G-CBM3a.

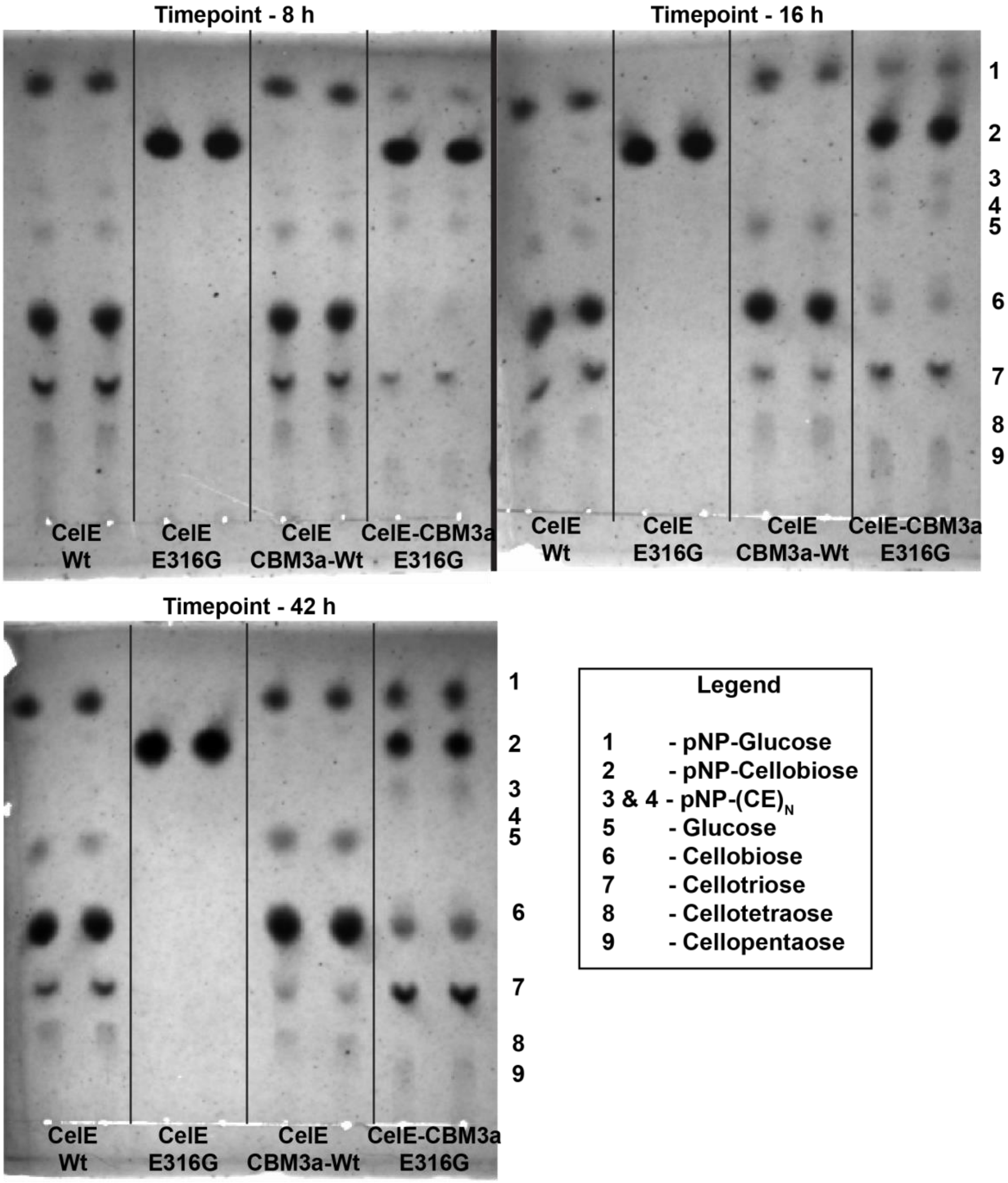


**Fig. S7. Representative TLC analysis after Orcinol staining for duplicate reaction mixtures of CelE, CelE-E316G, CelE-CBM3a-Wt, and CelE-CBM3a-E316G with pNP-cellobiose as starting substrate for varying reaction times.** Reaction conditions are identical to those reported in Figure 3A. Only limited time points (8 h, 16 h, 42 h) sampled from reactions conducted in Figure 3B are shown here for illustrative purposes to showcase the buildup of cello-oligosaccharide as products with increasing reaction time particularly for the CelE-CBM3a-E316G construct.

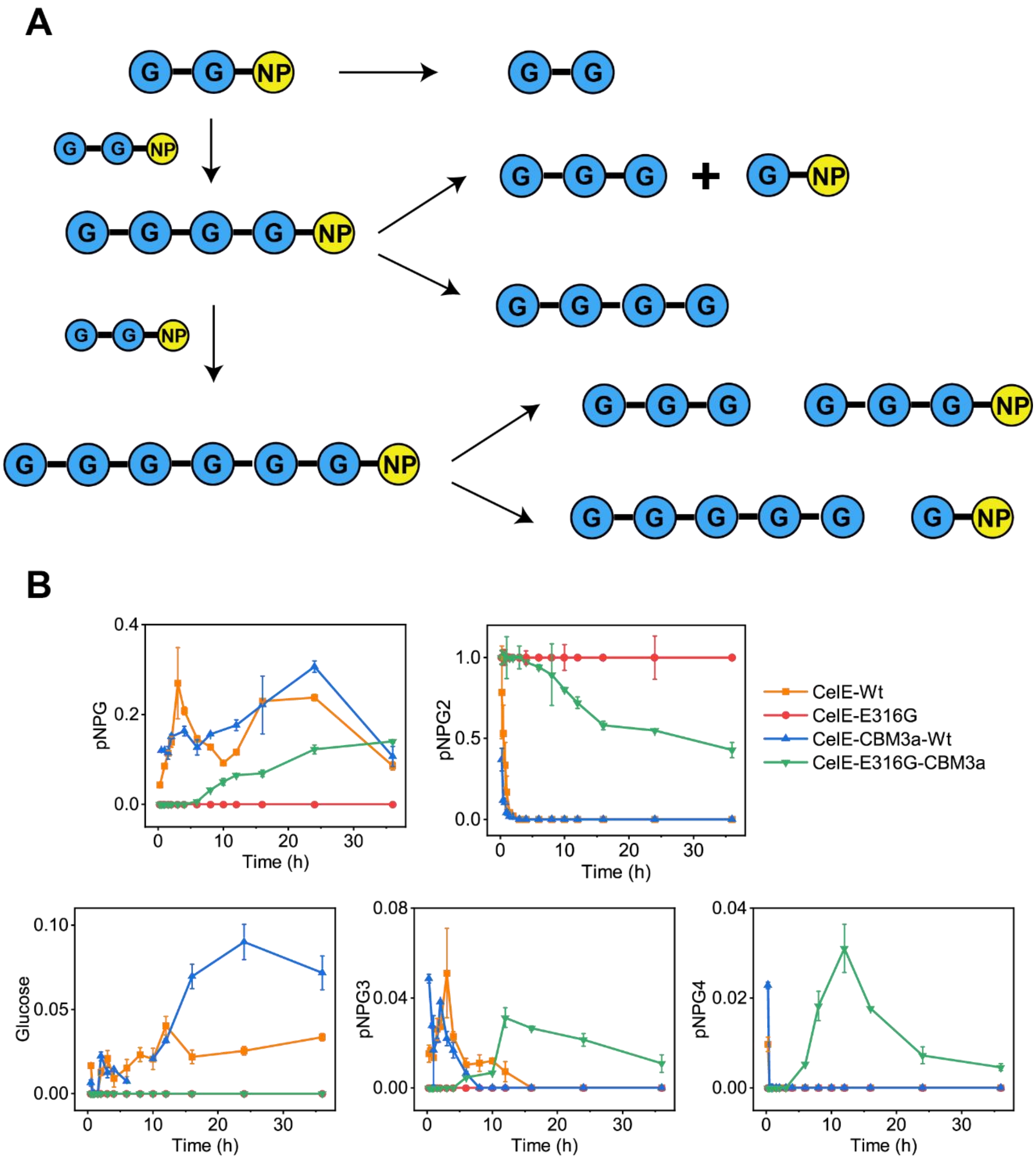


**Fig. S8. CelE-E316G-CBM3a utilizes a complex multiple-step reaction cascade, based on analyzed reaction products, to efficiently convert pNPC to mixture of complex glyco-products.** (A) Hypothesized reaction cascade scheme is shown here for all major reaction products observed with wild-type and E316G nucleophile mutants of CelE-CBM3a starting with pNPC as initial substrate. Glucose as product was only seen for wild-type enzymes (CelE-Wt and CelE-CBM3a-Wt) and is therefore not shown in cascade here. The blue circles indicate the glucosyl residue and yellow circle indicates the para-nitrophenyl group. (B) Detailed time course of fractional concentration profiles for various reaction products observed in the reaction mixtures as analyzed by quantitative thin layer chromatography (TLC) are shown here. Data is provided for CelE-Wt (orange square), CelE-CBM3a-Wt (blue triangle), CelE-E316G (red circle), and CelE-E316G-CBM3a (green inverted triangle) constructs. Error bar indicates one standard deviation from reported mean values from two biological replicates. Here, y-axis denotes fractional concentration of Glucose, pNPG: pNP-glucose, pNPG2: pNP-cellobiose, pNPG3: pNP-cellotriose, and pNPG4: pNP-cellotetraose. All components were estimated as pNPG2 (or pNPC) equivalent using pNPG2 as standard.

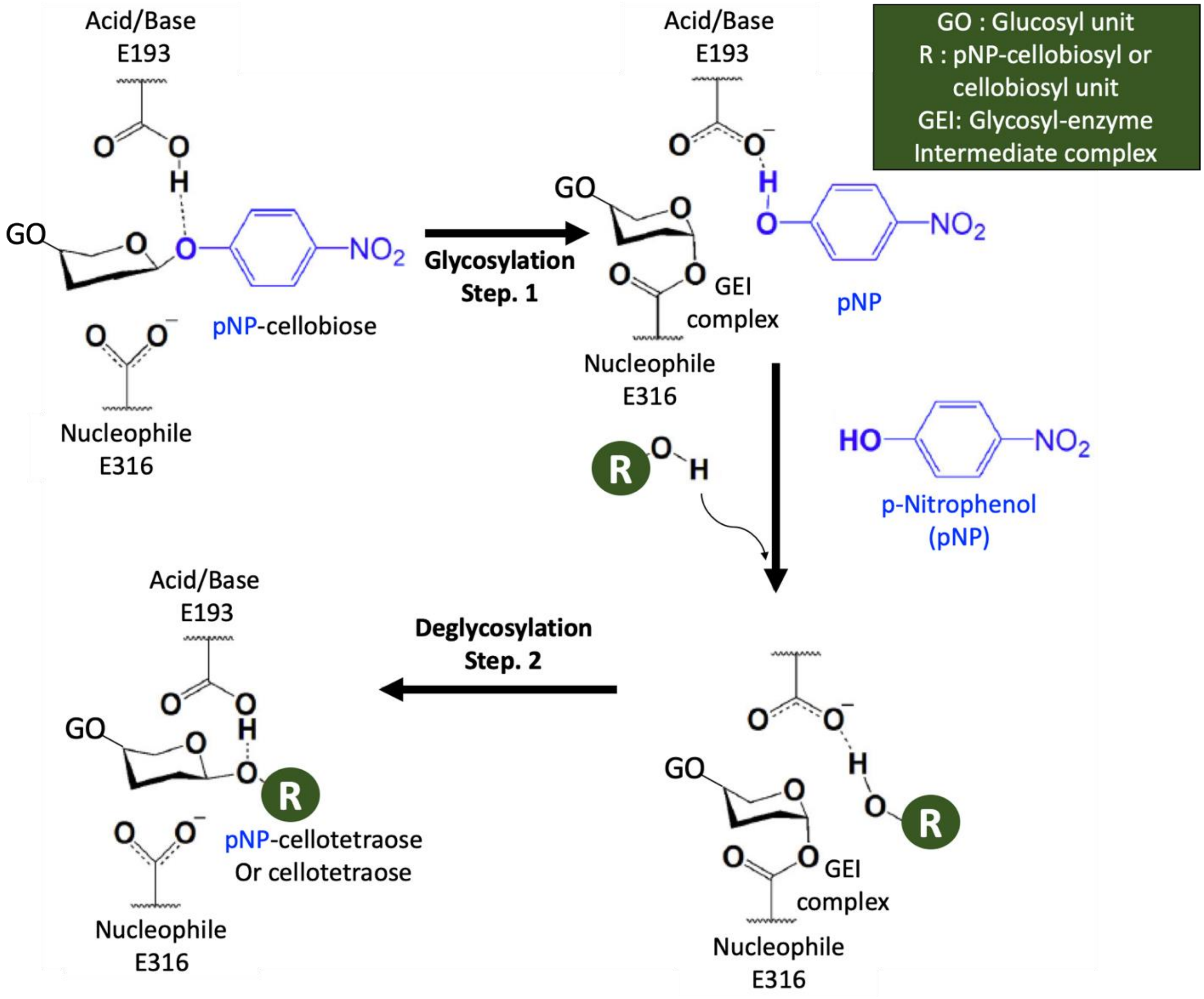


**Fig. S9. Glycosylation and deglycosylation steps via a conventional $S_N2$-type mechanism expected to operate for retaining β-retaining glycoside hydrolases like wild-type CelE is shown here.** Hydrolysis or transglycosylation reaction will take place if the acceptor moiety is water or another sugar group, respectively. Here, the nucleophile and acid/base residues are E316 and E193 for CelE, respectively, while the donor substrate is pNP-cellobiose and the acceptor substrate for transglycosylation can be either pNP-cellobiose (added substrate) or cellobiose (hydrolysis product). The final corresponding transglycosylation product for CelE is therefore expected to be either pNP-cellotetraose or cellotetraose, depending on the acceptor sugar group. Additional transglycosylation products can be additionally formed from these starting intermediates in a combinatorial manner. However, for native CelE as the reaction goes towards equilibrium, the transglycosylation products are eventually hydrolyzed into cellobiose mostly.

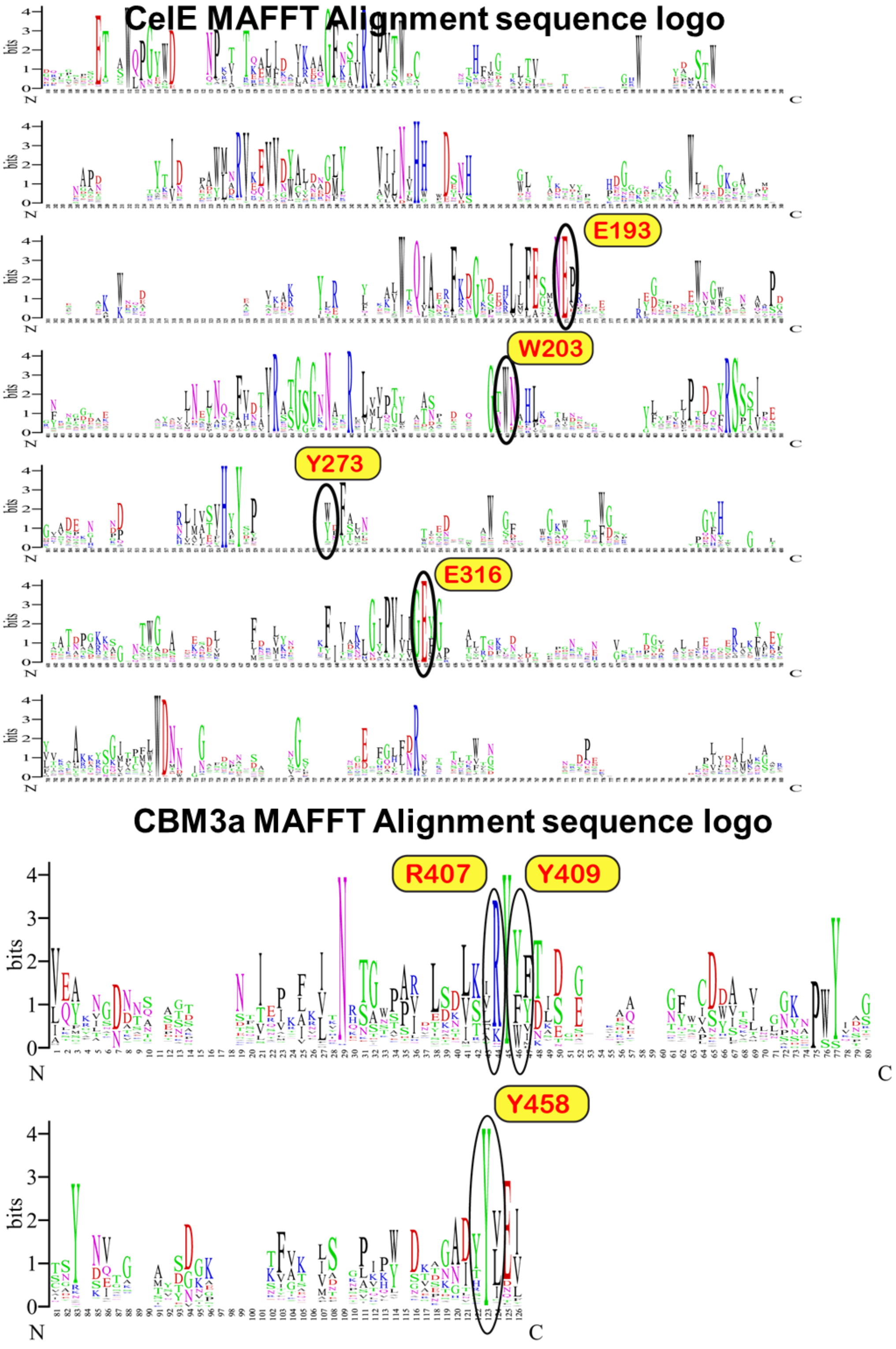


**Fig. S10. MAFFT alignments of CelE and CBM3a domains using Pfam database.** The sequence logo was generated using Geneious R11 software and several major conserved residues relevant to this study are highlighted here.

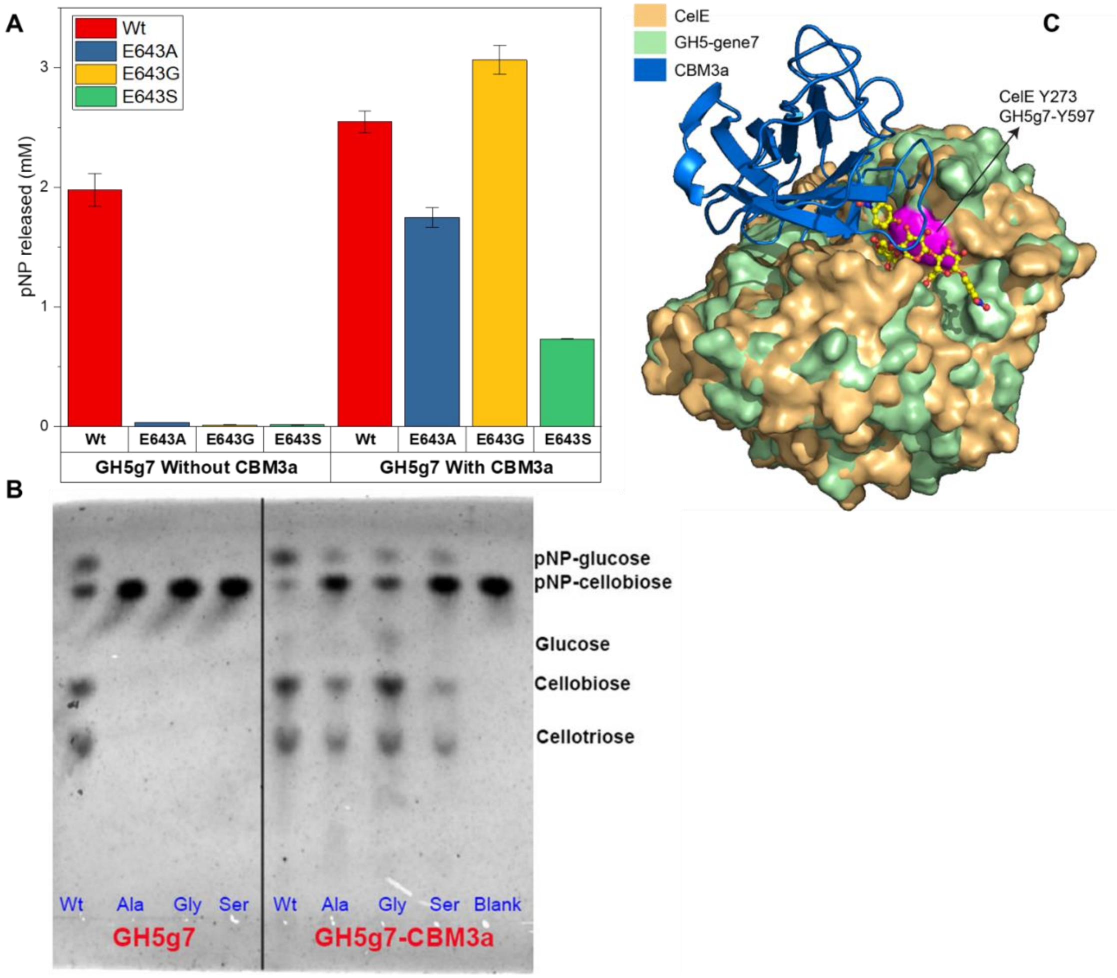


**Fig. S11. Activity of homologous GH5 family protein (GH5-gene7) on pNP-cellobiose.** GH5-gene7 belongs to the same GH5_4 subfamily as CelE enzyme (3). Here, 300 pmoles of the purified GH5g7 enzyme, with or without tethered CBM3a (analogous to CelE-CBM3a constructs), were reacted with 0.8 µmoles of pNP-cellobiose (pNP-CB) for 24 h at 40°C reaction temperature. All reactions were run in triplicates with error bars representing 1$\sigma$ from reported mean value. The relative amounts of pNP released (A) and TLC analysis by Orcinol staining (B) to evaluate the products formed in each reaction mixture are shown below. The 3D homologous structure of GH5-gene7 was docked along with the CelE structure to highlight how closely the two GH5_4 catalytic domains overlap structurally (C). CBM3a is docked adjacent to the CelE catalytic domain based on EOM analysis of SAXS data.

**Table S1.** Mutagenic forward (FP) and reverse (RP) primer sequences for performing site directed mutagenesis (A) and SLIC (B) to generate all constructs reported in this paper.

### A. Site-directed mutagenesis primers

| Name of Primer | Sequence |
|---|---|
| CelE_E316A_FP | GCTGTAATTATCGGAGCATTCGGAACCATTGAC |
| CelE_E316A_RP | GTCAATGGTTCCGAATGCTCCGATAATTACAGC |
| CelE_E316G_FP | GCTGTAATTATCGGAGGATTCGGAACCATTGAC |
| CelE_E316G_RP | GTCAATGGTTCCGAATCCTCCGATAATTACAGC |
| CelE_E316S_FP | GCTGTAATTATCGGATCATTCGGAACCATTGAC |
| CelE_E316S_RP | GTCAATGGTTCCGAATGATCCGATAATTACAGC |
| CelE_E193A_FP | GAGACAATGAACGCACCGAGAGAAG |
| CelE_E193A_RP | CTTCTCTCGGTGCGTTCATTGTCTC |
| CelE_Y273A_FP | GCTTATTCACCGGCTTTCTTTGCTATGG |
| CelE_Y273A_RP | CCATAGCAAAGAAAGCCGGTGAATAAGC |
| CelE_Y270A_FP | GTATCCATACATGCTGCTTCACCGTATTTCTTTG |
| CelE_Y270A_RP | CAAAGAAATACGGTGAAGCAGCATGTATGGATAC |
| CelE_W203A_FP | TAGGTTCACCTATGGAAGCAATGGGCGGAACGTATG |
| CelE_W203A_RP | CATACGTTCCGCCCATTGCTTCCATAGGTGAACCTA |
| CelE_H268A_FP | AGTAATAGTATCCATAGCTGCTTATTCACCGTAT |
| CelE_H268A_RP | ATACGGTGAATAAGCAGCTATGGATACTATTACT |
| CelE_W349A_FP | GAATTGCTGTTTTCTGGGCTGATAACGGCTATTAC |
| CelE_W349A_RP | GTAATAGCCGTTATCAGCCCAGAAAACAGCAATTC |
| CBM3a_Y458A_FP | CAATGCTGATACCGCCCTGGAAATTAGC |
| CBM3a_Y458A_RP | GCTAATTTCCAGGGCGGTATCAGCATTG |
| CBM3a_E521A_FP | GTTTGGGGGAAAGCACCAGGATAGTAG |
| CBM3a_E521A_RP | CTACTATCCTGGTGCTTTCCCCCAAAC |
| CBM3a_R407A_FP | CGAAACTGACCCTTGCTTACTACTATACGG |
| CBM3a_R407A_RP | CCGTATAGTAGTAAGCAAGGGTCAGTTTCG |
| CBM3a_T509A_FP | GGGATCAGGTGGCCGCATATTTGAAC |
| CBM3a_T509A_RP | GTTCAAATATGCGGCCACCTGATCCC |
| CBM3a_Y409A_FP | GACCCTTCGTTACGCCTATACGGTTGATG |
| CBM3a_Y409A_RP | CATCAACCGTATAGGCGTAACGAAGGGTC |

### B. Sequence and ligation independent cloning (SLIC) primers

| Name of Primer | Sequence |
|---|---|
| 6aa_Linker_FP | GACTCCCACTAAAGTAAGCGGTAACCTGAAGGTTG |
| 6aa_Linker_RP | GGTTACCGCTTACTTTAGTGGGAGTCGCGTTTAAAC |
| 11aa_Linker_FP | GTGCCACTCCTACCGTAAGCGGTAACCTGAAGGTTG |
| 11aa_Linker_RP | GGTTACCGCTTACGGTAGGAGTGGCACCTTTAGTG |
| 21aa_Linker_FP | CTAAGTCGGCAACGGTAAGCGGTAACCTGAAGGTTG |
| 21aa_Linker_RP | GGTTACCGCTTACCGTTGCCGACTTAGTCGGAG |
| Flex1_Vec_FP | GCGAACACCGGAGTAAGCGGTAACCTGAAGG |
| Flex1_Vec_RP | GCCAGTCGCGTTTAAACCTTCAACGCCGGC |
| Flex1_Ins_FP | GTTGAAGGTTTAAACGCGACTGGCACTAAAGGTG |
| Flex1_Ins_RP | GTTACCGCTTACTCCGGTGTTCGCCCCGGTA |

| | |
|---|---|
| Flex2_Vec_FP | CGAACGGAGGAGTAAGCGGTAACCTGAAGG |
| Flex2_Vec_RP | GCCACTCGCGTTTAAACCTTCAACGCCGGC |
| Flex2_Ins_FP | TTGAAGGTTTAAACGCGAGTGGCGGTAAAG |
| Flex2_Ins_RP | GTTACCGCTTACTCCTCCGTTCGCCCCTCC |
| Rig1_Vec_FP | GCGAACACCCCAGTAAGCGGTAACCTGAAGGTTG |
| Rig1_Vec_RP | GGGAGTCGCGTTTAAACCTTCAACGCCGGC |
| Rig1_Ins_FP | GTTGAAGGTTTAAACGCGACTCCCACTAAACCTG |
| Rig1_Ins_RP | TTACCGCTTACTGGGGTGTTCGCCGGGG |
| Rig2_Vec_FP | CGAGCGGTGGCGTAAGCGGTAACCTGAAGG |
| Rig2_Vec_RP | GCGCTGCCACCTAAACCTTCAACGCCGGC |
| Rig2_Ins_FP | TTGAAGGTTTAGGTGGCAGCGCGGAGGCT |
| Rig2_Ins_RP | GTTACCGCTTACGCCACCGCTCGCCTTCGC |

**Table S2.** Real space radius of gyration ($R_g$) and $D_{max}$ for CelE-E316G and CelE-E316G-CBM3a protein constructs, with or without added increasing pNP-cellobiose substrate concentrations (i.e., 0, 258, 644 moles pNPC per mole protein), were estimated using the p(r) function fitted to the SAXS data.

| **Sample Type** | $R_g$ **(Å)** | $D_{max}$ **(Å)** |
|---|---|---|
| His-CD (i.e., CelE-E316G) | 28 ± 1 | 102 |
| His-CD-Linker-CBM3a (No substrate) | 44 ± 1 | 154 |
| His-CD-Linker-CBM3a: Substrate (1:258) | 43 ± 2 | 155 |
| His-CD-Linker-CBM3a: Substrate (1:644) | 45 ± 1 | 157 |
